# Formation of functional, non-amyloidogenic fibres by recombinant *Bacillus subtilis* TasA

**DOI:** 10.1101/188995

**Authors:** Elliot Erskine, Ryan J Morris, Marieke Schor, Chris Earl, Rachel M. C. Gillespie, Keith Bromley, Tetyana Sukhodub, Lauren Clark, Paul K. Fyfe, Louise C. Serpell, Nicola R. Stanley-Wall, Cait E. MacPhee

## Abstract

Bacterial biofilms are communities of microbial cells encased within a self-produced polymeric matrix. In the *Bacillus subtilis* biofilm matrix the extracellular fibres of TasA are essential. Here a recombinant expression system allows interrogation of TasA, revealing that monomeric and fibre forms of TasA have identical secondary structure, suggesting that fibrous TasA is a linear assembly of globular units. Recombinant TasA fibres form spontaneously, and share the biological activity of TasA fibres extracted from *B. subtilis*, whereas a TasA variant restricted to a monomeric form is inactive and subjected to extracellular proteolysis. The biophysical properties of both native and recombinant TasA fibres indicate that they are not functional amyloid-like fibres. A gel formed by TasA fibres can recover after physical shear force, suggesting that the biofilm matrix is not static and that these properties may enable *B. subtilis* to remodel its local environment in response to external cues. Using recombinant fibres formed by TasA orthologues we uncover species variability in the ability of heterologous fibres to cross-complement the *B. subtilis tasA* deletion. These findings are indicative of specificity in the biophysical requirements of the TasA fibres across different species and/or reflect the precise molecular interactions needed for biofilm matrix assembly.

**Contributions:** Conceived and designed the experiments: CE, EE, RG, CEM, RJM, MS, NSW; Performed the experiments: KB, LC, CE, EE, PKF, RG, CEM, RJM, MS, TS; Contributed new analytical tools: CE, EE, RG, TS; Analysed the data: CE, EE, CEM, RJM, MS, LCS, NSW; Wrote the paper: EE, RJM, CEM, MS, NSW.

## Introduction

Biofilms are communities of microbial cells that underpin diverse processes including sewage bioremediation, plant growth promotion, chronic infections and industrial biofouling (Costerton *et al*., 1987). The microbial cells resident in the biofilm are encased within a self-produced extracellular polymeric matrix that commonly comprises lipids, proteins, extracellular DNA, and exopolysaccharides (Flemming and Wingender, 2010; Hobley *et al*., 2015). This matrix fulfils a variety of functions for the community, from providing structural rigidity and protection from the external environment, to supporting signal transduction and nutrient adsorption (Flemming and Wingender, 2010; Dragoš and Kovács, 2017; Vidakovic *et al*., 2018). *Bacillus subtilis* is a soil dwelling bacterium that is a model for biofilm formation by Gram-positive bacteria; beyond this it is of commercial interest due to its biocontrol and plant growth promoting properties that highlight its potential to substitute for petrochemical derived pesticides and fertilizers (Bais *et al*., 2004; Chen *et al*., 2012; Chen *et al*., 2013). Biofilm formation is subject to complex regulatory pathways (Cairns *et al*., 2014) and it is known that the *B. subtilis* biofilm matrix predominantly comprises three specific components. The first is an exopolysaccharide that serves to retain moisture within the biofilm and functions as a signalling molecule (Seminara *et al*., 2012; Elsholz *et al*., 2014). The composition of the exopolysaccharide remains unclear due to three inconsistent monosaccharide composition analyses being detailed thus far (Chai *et al*., 2012; Jones *et al*., 2014; Roux *et al*., 2015). The second component is the protein BslA that is responsible for the non-wetting nature of the biofilm (14, 15). We have previously determined the structure of BslA (Hobley *et al*., 2013), identified a unique structural metamorphosis that enables BslA to self-assemble at an interface in an environmentally-responsive fashion (Bromley *et al*., 2015), and discovered that BslA is also important for determining biofilm architecture, independently of its ability to render the surface of the biofilm water-repellent (Arnaouteli *et al*., 2017). The third component of the biofilm matrix is the protein TasA (together with accessory protein TapA) that is needed for biofilm structure including attachment to plant roots (Branda *et al*., 2004; Romero *et al*., 2011; Beauregard *et al*., 2013).

TasA is a product of the *tapA-sipW-tasA* locus (Michna *et al*., 2016). It is post-translationally modified by SipW (Stöver and Driks, 1999), a specialized signal peptidase that releases the mature 261-amino acid TasA into the extracellular environment where it forms long protein fibres that contribute to the superstructure of the biofilm matrix and are needed for biofilm integrity (Branda *et al*., 2006; Romero *et al*., 2010). In addition to functions involved in the process of biofilm formation, TasA is also linked with sliding motility (van Gestel *et al*., 2015) and spore coat formation (Stöver and Driks, 1999; Serrano *et al*., 1999). TasA fibres can be extracted from *B. subtilis* biofilms, and exogenous provision to a *tasA* null strain has previously been reported to reinstate structure to floating pellicles (Romero *et al*., 2010). Due to the reported ability of TasA fibres to bind the dyes Congo Red and Thioflavin T (ThT), *ex vivo* purified TasA fibres have previously been classified as functional bacterial amyloid fibres (Romero *et al*., 2010), placing them alongside the curli fibres of *E. coli* (Chapman *et al*., 2002).

Amyloid-like fibres are well-known for their association with diseases like Alzheimer’s and Parkinson’s (Eisenberg and Jucker, 2012). In these conditions, highly stable fibrillar protein deposits are found in tissue sections, and are associated with cell damage (Hardy and Selkoe, 2002). The amyloid fibres in these deposits are characterised by several properties: i) β-sheet-rich structures that are assembled into the canonical “cross-β” structure; ii) the ability of the fibres to bind the dye Congo Red and exhibit green birefringence under cross polarised light; iii) kinetics of formation that indicate a nucleated self-assembly process; and iv) a fibril structure that is unbranched, 6-12 nm in diameter, and often microns in length (Sunde and Blake, 1997; Sipe *et al*., 2016). Once formed, these protein aggregates are highly stable, and in many cases are thought to be the lowest energy structural form shorter polypeptide chains can adopt (Baldwin *et al*., 2011). ‘Functional’ amyloid fibres refer to fibrillar protein deposits that share the characteristic structural properties of amyloid fibres, but are beneficial to the organism rather than being associated with disease (Fowler *et al*., 2007). Significant caution is required in identifying functional amyloid-like fibres from predominantly *in vitro* data however, as many proteins and peptides can be induced to adopt the canonical amyloid fibre cross-β fold through appropriate manipulation of solution conditions such as changes in pH, temperature, cosolvent, salt or the presence of an interface (e.g. 34–38), which may or may not be of physiological relevance. Indeed, the ability of proteins to assemble into the cross-β architecture appears to be a ‘generic’ property of the polypeptide chain, independent of the amino acid sequence or the native structure of the precursor (Dobson, 1999; MacPhee and Dobson, 2000).

Here we show that, although TasA is a fibre-forming protein, it is not amyloid-like in character. We have produced recombinant TasA in both fibre and monomeric forms, and show that the secondary structures of these are both identical to each other and to those reported previously for the exogenous purified TasA fibres (Romero *et al*., 2010; Chai *et al*., 2013), appearing significantly helical in character. We have also examined native TasA fibres in enriched extracts from *B. subtilis* and show that both the native and recombinant forms of fibrous TasA show indistinguishable biological activity, being able to reinstate biofilm structure to a Δ*tasA sinR* deletion strain. X-ray fibre diffraction of the recombinant TasA fibres shows that they are assembled from a helical repeat of globular protein units arranged approximately 45 Å apart, and the data are not consistent with the canonical “cross-β” diffraction pattern associated with amyloid-like fibres. Neither monomeric nor fibrous forms of recombinant TasA bind the dyes Congo Red or ThT, and although TasA-enriched extracts from *B. subtilis* biofilms show both Congo Red and ThT binding activity, this is at a similar level to that produced by protein extracts from cells lacking *tasA*. Thus, TasA does not fall into the class of “functional amyloid-like fibres”; nonetheless it plays a critical role in biofilm structure.

## Results

### TasA forms non-amyloid fibres and is rendered monomeric by placement of a single N-terminal amino acid

We predicted the identity of the N-terminus of mature TasA protein *in silico* using SignalP v4.2 (Petersen *et al*., 2011) and subsequently confirmed this *in vivo* using mass spectrometry. Based on this information we designed an expression construct to allow purification of recombinant *B. subtilis* TasA (Fig. S1A), corresponding to the mature TasA sequence covering amino acids 28-261, after production in *E. coli* (Fig. S1B). The purified protein displayed obvious viscosity, not flowing upon inversion of the tube, and bead tracking microrheology confirmed the gel-like nature of the solution (Fig. S1C). This viscosity arises from the formation of a fibrous aggregate that can be characterised by transmission electron microscopy (TEM) (Fig. 1A). Within these fibres we observed a subunit repeat along the fibre axis, repeating at approximately 4 nm. Hereafter we refer to this protein as ‘fTasA’. To compare the recombinant protein to the native form, we extracted TasA fibres from *B. subtilis*, which we refer to as nTasA(+). To identify the specific contribution of TasA originating from this partially purified, heterogeneous sample we followed the same enrichment process with a strain carrying a *tasA* deletion, this sample is referred to as nTasA(−) (Fig. S1D-E). The subunit repeat pattern seen in the recombinant TasA fibres was also visible in the nTasA(+) fibres (Fig 1B) and no comparable fibres were observed in the nTasA(−) sample (Fig. S1F).

**Figure 1:**
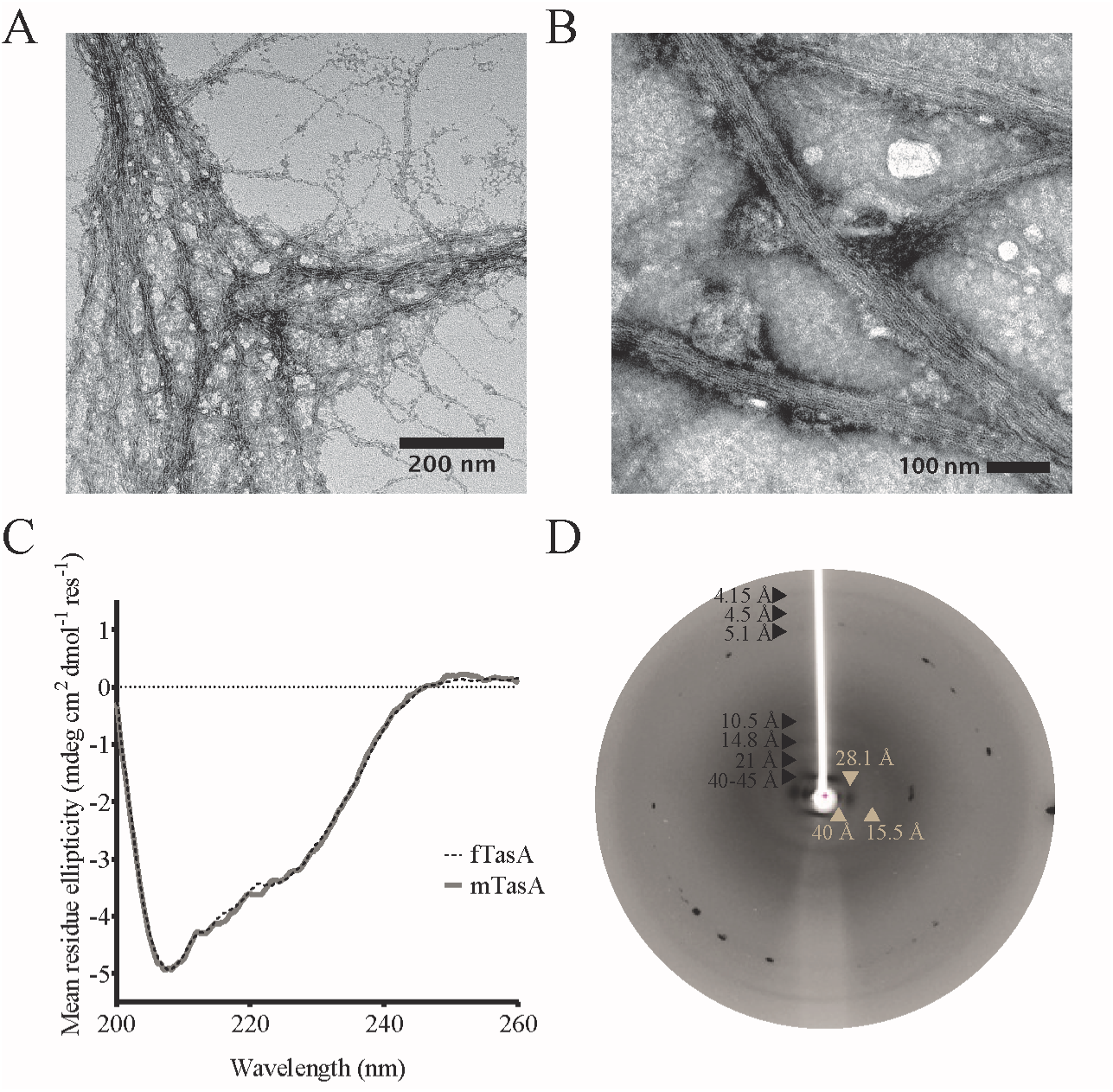
Recombinant TasA forms fibres. (**A-B**) Transmission electron microscopy images of recombinant fTasA and nTasA(+) stained with uranyl acetate shows the presence of fibres several microns in length and approximately 15 nm wide (**C**) Solution state circular dichroism spectra of recombinant fTasA (Dotted black line) and mTasA (Solid grey line). (**D**) X-Ray Diffraction of recombinant fTasA protein fibres with exposure for 60 seconds where meridional and equatorial diffraction signals indicated in black and beige respectively.

Circular dichroism (CD) spectroscopy of fTasA shows a minimum at 208 nm, a shoulder at ~222 nm, and a maximum below 200 nm (Fig. 1C). The overall shape and the position of the minima are consistent with a predominantly (>50%) α-helical conformation, however the ratio of the two minima suggests there are likely to be contributions from other secondary structural elements. The spectrum resembles that previously reported for TasA oligomers (Chai *et al*., 2013) or fibres purified directly from *B. subtilis* (Romero *et al*., 2010), and does not display the high β-sheet content (represented by a single minimum at ~216-218 nm) typical for proteins in amyloid-like fibres; nor do the measured minima/maximum correspond to those predicted or measured for highly twisted β-sheets (Micsonai *et al*., n.d.). It was not possible to obtain a CD spectrum for the natively extracted nTasA(+) due to contamination with flagella (and other proteins), which were also visible in TEM images (Fig. S1F, arrow), and identified due to their uniformity and their presence in the nTasA(−) extract.

X-ray fibre diffraction from partially aligned fTasA fibres (Fig. 1D) showed a series of layer lines on the meridian. The lowest resolution visible was 41-45 Å/4.1-4.5 nm and further strong layer lines were measured at 21 Å and 14.8 Å. A weak, high resolution meridional diffraction signal was observed at 4.15 Å. These spacings are consistent with a helical or globular repeat distance of 40-45 Å, as was observed by TEM for both fTasA and nTasA(+). On the equator of the pattern, strong diffraction signals were measured at approx. 40 Å, 28.1 Å and 15.5 Å. These spacings are consistent with packing of a fibre with globular units of dimensions 28 × 45 Å. The diffraction signals expected for a cross-β amyloid-like structure (i.e. 4.7 Å (the inter-strand distance) and ~10-12 Å (the inter-sheet distance) (O Sumner Makin and Serpell, 2005)) were not observed.

Many amyloid-like fibres bind the dyes Congo Red and ThT and dye-binding assays are often used to assess fibril assembly, so we next tested whether our recombinant protein fTasA and our *B. subtilis* nTasA(+) extract bound these dyes. ThT fluorescence in the presence of fTasA was similar to that of a non-fibrillar control protein (Fig. S1G), and showed no evidence of an interaction with Congo Red (Fig. S1H; fibrils assembled from insulin are shown as a positive control). The nTasA(+) extract enhanced ThT fluorescence and showed Congo Red binding (Fig. S1G-H), however so did the nTasA(−) extract from the strain carrying the *tasA* deletion (Fig. S1G-H). Therefore, there is no evidence that fibre-forming TasA binds ThT or Congo Red, whether recombinant or extracted from *B. subtilis*. Thus, the combination of the visible subunit repeat in both recombinant and native TasA fibres, the absence of a clear β-sheet secondary structure in the recombinant protein (as has also been reported for *ex vivo* TasA), the X-ray fibre diffraction results, and the lack of dye binding, all suggest that TasA is not a functional amyloid fibre (see *Discussion*).

Through addition of a single amino acid to the N-terminus of the mature TasA protein (Fig. S2A), we discovered that it was possible to block fibre formation *in vitro*. Inhibition of TasA fibre assembly was not dependent on the chemical properties of the added amino acid, with lysine, phenylalanine, glutamic acid, alanine, or serine all being effective (Fig. S2B). Having compared the behaviour of these proteins at this level we focussed our subsequent analysis on the purified monomeric serine-tagged TasA (Fig. S2C). Size-exclusion chromatography (SEC) showed a single peak of around 30 kDa (Fig. S2D), and the molecular mass was confirmed by LC-MS, ensuring the specificity of the protein sequence. Monomeric TasA (‘mTasA’) was not viscous, with bead tracking microrheology confirming the liquid-like nature of the purified protein solution (Fig. S1C); moreover, no fibres were apparent by TEM. The monomeric protein showed the same lack of ThT binding as fTasA (Fig. S1G). The CD spectrum of mTasA was indistinguishable from fTasA (Fig. 1C) indicating that addition of a single amino acid to the N-terminus did not affect the secondary structure, and moreover suggesting that the fibrous form is likely constructed from a linear assembly of these monomeric units. Thus, examination of TasA form and function in both fibrous and monomeric states was possible.

### Recombinant TasA fibres are biologically active

To test the biological functionality of the recombinant TasA fibres, an in frame Δ*tasA* deletion (NRS5267) was constructed. Exogenous addition of neither recombinant fTasA (Fig. S2A), nor purified nTasA(+) successfully reinstated biofilm architecture to the Δ*tasA* mutant (Fig. 2A, S3B). Immunoblot analysis of whole biofilm protein extracts using anti-TasA antibodies revealed that while fTasA and nTasA(+) do not influence biofilm morphology, when cultured with the *tasA* deletion strain, the exogenously added proteins are still detectable after 48 hours incubation (Fig. 2B, S3C). The biofilm phenotype was similarly unchanged when monomeric TasA or nTasA(−) was exogenously added to the Δ*tasA* strain (Fig. 2A, S3B). For the monomeric protein, we considered two possibilities: i) mTasA is unstable in the presence of cells when added exogenously, or ii) mTasA is stable, but not functional, suggesting that it is not converted into a functional form following exogenous addition to the biofilm. Immunoblot analysis of protein extracts using anti-TasA antibodies revealed that exogenously added mTasA reached undetectable levels after incubation with the *tasA* strain during biofilm formation conditions (Fig. 2B). These findings indicate that mTasA is likely to be degraded by proteolysis.

**Figure 2:**
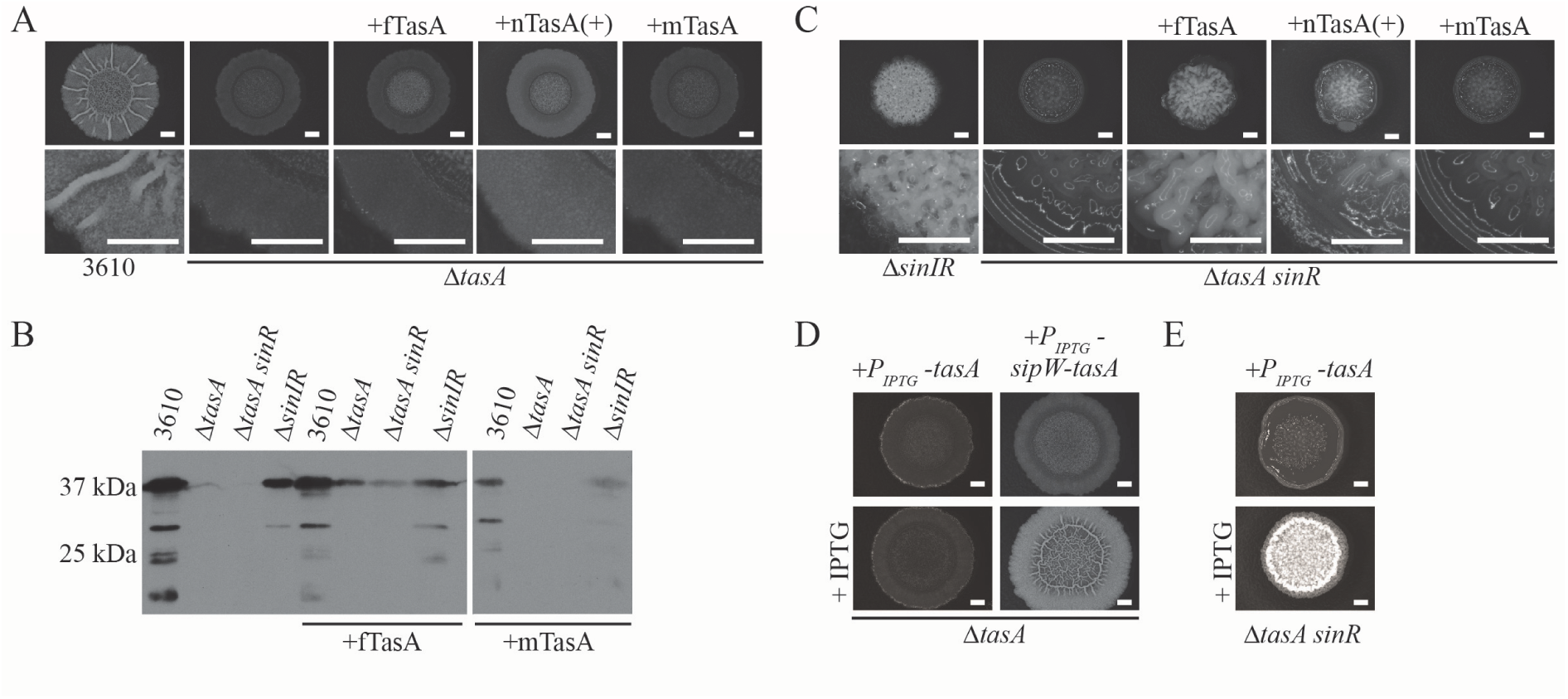
Recombinant fTasA is biologically active. (**A,C**) Biofilm phenotypes of wild type (NCIB3610), Δ*tasA* (NRS5267), Δ*sinIR* (NRS2012) and Δ*tasA sinR* (NRS5248) strains with the addition of 10 μg fTasA, 30 μg nTasA(+) or 10 μg mTasA as indicated. (**B**) Immunoblot blot analysis of biofilm lysate collected from biofilms challenged with α-TasA antibody. (**D-E**) Biofilm phenotype of ΔtasA and Δ*tasA sinR* complementation in presence of 100 μM for Δ*tasA* P_iptg_-*tasA* (NRS5276) and 25 μM for Δ*tasA sinR* P_iptg_-*tasA* (NRS5255) and Δ*tasA* P_iptg_-*sipW-tasA* (NRS5313).

*B. subtilis* secretes 7 heat-labile proteases into the extracellular environment (Sloma *et al*., 1988; Rufo *et al*., 1990; Sloma *et al*., 1990; Wu *et al*., 1990; Tran *et al*., 1991; Sloma *et al*., 1991; Margot and Karamata, 1996) to which mTasA could be exposed. Therefore we analysed the stability of recombinant fTasA and mTasA after incubation in cell-free spent culture supernatant derived from planktonic growth of NCIB3610 to stationary phase. This revealed that assembly of TasA into the fibre form confers protection from degradation (Fig. S3D-E). We then assessed stability of mTasA and fTasA protein in spent culture supernatant isolated from a laboratory prototrophic strain (PY79) and two derivatives of PY79 that lacked the coding regions for secreted extracellular proteases: namely strains in which the genes for six (“Δ6”) or seven (“Δ7”) of the native proteases had been deleted (Table S1). While fTasA was detected under all incubation conditions (Fig. S3D), mTasA was only observed when the culture supernatant had been heat treated to denature the protein content or when it was incubated with the spent culture supernatant derived from Δ6 and Δ7 exoprotease-deficient strains (Fig. S3E). These results indicate susceptibility of the monomeric TasA protein to proteolysis and that protection is conferred by self-assembly to a fibre form.

SinR is a major repressor of biofilm formation that functions, in part, by directly inhibiting transcription from the operons needed for the production of the exopolysaccharide and TasA fibres, both essential components of the *B. subtilis* matrix (Chu *et al*., 2006). Deletion of *sinR* results in a biofilm that is densely wrinkled and highly adherent to a surface when compared to the parental strain, due to increased production of the biofilm macromolecules (Fig. 2C, S3B). While constructing the Δ*tasA* strain we inadvertently isolated a Δ*tasA sinR* deletion strain (Fig. 2C, S3B) which displayed a flat, featureless biofilm by comparison with a *sinR* deletion strain. Serendipitously, we found that addition of 10 μg recombinant fTasA or 30 μg of nTasA(+) extract broadly returned the wrinkled *sinR* mutant-like phenotype to the Δ*tasA sinR* mutant (Fig. 2C, S3B). Thus, the recombinant form of fTasA is biologically functional, and shows the same functional activity as native TasA. In contrast, monomeric TasA (Fig. S2A) and the nTasA(−) samples did not reinstate rugosity to the Δ*tasA sinR* deletion strain (Fig. 2C, S3B) suggesting that *in vivo* templating of mTasA into a functional fibrous form does not occur and that the activity in the nTasA(+) sample was linked to TasA activity specifically. As was observed when protein was supplied exogenously to the single *tasA* deletion strain, mTasA was not detectable by anti-TasA immunoblot analysis after co-incubation with the Δ*tasA sinR* deletion strain, but fTasA was detected (Fig. 2B), confirming the susceptibility of mTasA to proteolysis.

We cannot explain why fTasA and nTasA(+) do not recover biofilm rugosity to the *tasA* deletion when supplied exogenously. However we note that the *tasA* and *tasA sinR* strains differ in the requirements needed for genetic complementation. The *tasA* deletion cannot be genetically complemented by expression of *tasA* under the control of an inducible promoter at the ectopic *amyE* locus (Fig. 2D, S3F) but requires co-expression of *sipW* and *tasA* to return biofilm formation to a wild-type morphology (Fig. 2D, S3F). This is not an indication that *sipW* is inadvertently disrupted in the *tasA* strain, as restoration of biofilm formation by the *tasA* mutant was equally successful using a complementation construct when codons 3 and 4 of *sipW* were replaced with stop codons (Fig. S3F). In contrast, provision of the *tasA* coding region only at the ectopic *amyE* locus in the Δ*tasA sinR* deletion strain (NRS5255) is sufficient to allow a densely wrinkled biofilm structure to be recovered (Fig 2E, S3G). We next explored the mechanism underpinning the interaction between fibrous TasA and the components of the biofilm.

### Recombinant TasA fibres require the biofilm exopolysaccharide for activity

TasA protein fibres have been reported to be anchored to the cell wall *via* an interaction with a partner protein called TapA (Romero *et al*., 2011). Moreover, deletion of *tapA* is associated with a reduction in the level of TasA (Romero *et al*., 2014). As the deletion of *sinR* leads to increased transcription of the entire *tapA* operon (Chu *et al*., 2006), we hypothesised that there may be an increase in available TapA ‘docking’ sites available for the anchoring of TasA fibres when added *ex vivo* to the Δ*tasA sinR* double mutant. To test if TapA is needed for wrinkling of the Δ*tasA sinR* deletion strain upon addition of preassembled TasA fibres, we constructed a Δ*tapA* Δ*tasA sinR* triple deletion strain (Fig. 3A, S4A). This strain could be returned to the *sinR* morphology upon genetic complementation with the *tapA-sipW-tasA* gene cluster at an ectopic location in the chromosome (Fig. S4B). When fTasA was co-cultured with the Δ*tapA* Δ*tasA sinR* strain, we observed similar levels of *ex vivo* complementation as when fTasA was added to the Δ*tasA sinR* deletion strain (compare Fig. 2C and 3A), thus suggesting that TapA is not required to reinstate biofilm architecture when fully formed TasA fibres are supplied.

**Figure 3:**
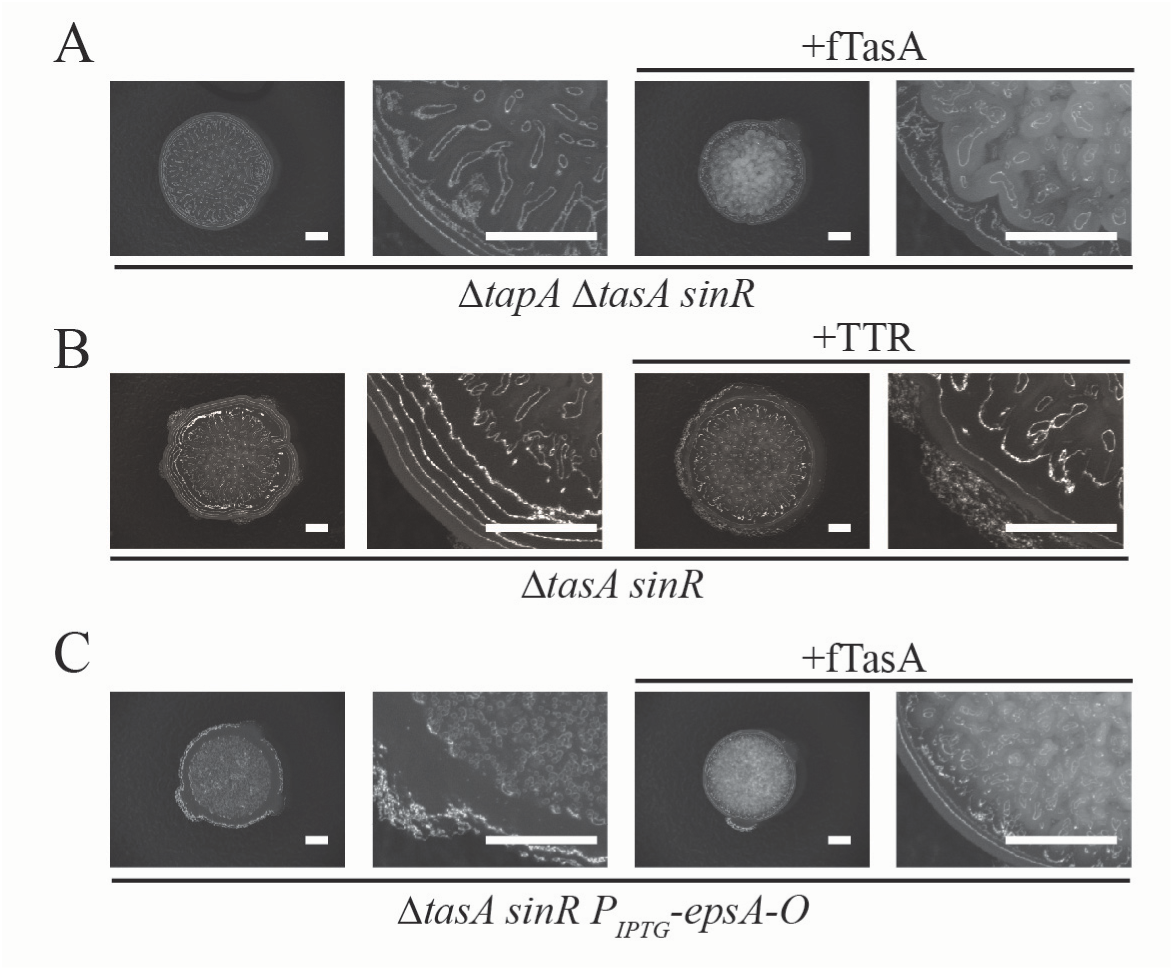
Recombinant fTasA does not need TapA but required the biofilm exopolysaccharide for activity. **(A)** Biofilm phenotype of Δ*tapA* Δ*tasA sinR* (NRS5749) mutant upon addition of 10 μg fTasA *ex vivo*. (**B**) Biofilm phenotype of Δ*tasA sinR* (NRS5248) upon addition of transthyretin (TTR). (**C**) Biofilm phenotype of Δ*tasA sinR* P_iptg_-*epsA-O* (NRS5421) strain in the presence of 100 μM IPTG in absence and presence of *ex vivo* addition of 10 μg fTasA. An enlarged section of bottom left corner of the biofilm is shown.

In light of the findings above we explored if the rugosity displayed by the Δ*tasA sinR* in the presence of *ex vivo* recombinant fTasA was due to a specific interaction with the matrix components, or whether the presence of sufficient fibrous material is enough to confer rugosity simply due to the gelatinous nature of the concentrated fTasA protein. To test this, we took two approaches. First we tested if an entirely unrelated protein fibre could substitute for fTasA, simply by provision of a fibrous protein scaffold. We provided amyloid-like fibres assembled from the well-characterised transthyretin peptide 105-115 (TTR_105-115_) (Fitzpatrick *et al*., 2013) exogenously to the Δ*tasA sinR* strain followed by incubation under biofilm forming conditions. Despite the obvious viscosity of the TTR_105-115_ gel, it did not reinstate biofilm rugosity (Fig. 3B, S4D). Therefore, a biochemically distinct fibre cannot substitute for fTasA. Next, we assessed whether the biofilm exopolysaccharide was needed for rugosity under these conditions. This experiment was based on the premise that if the wrinkle formation after addition of exogenous fTasA was derived from the gelling properties of fTasA, the exopolysaccharide would not be needed. To determine this we used a strain where the entire *epsA-O* operon was placed under the control of an IPTG inducible promoter at the native location on the chromosome (Terra *et al*., 2012). We then added fTasA with and without induction of the *epsA-O* operon. Analysis of the biofilm phenotypes revealed that we were able to induce rugosity with fTasA only in the presence of IPTG (Fig. 3C, S4E), although not to the same extent as seen in the parent strain - most likely because production of the exopolysaccharide is uncoupled from its native regulation circuitry, impacting the level of polymer produced. Therefore we can conclude that both the biofilm exopolysaccharide and TasA are required to return rugosity to the biofilm.

### Biophysical properties of recombinant orthologous TasA

Using the *B. subtilis* TasA protein sequence we identified orthologous proteins from a range of *Bacillus* species. The sequences were aligned using Clustal Omega (Sievers *et al*., 2011) (Fig. S5) and used to generate a phylogenetic tree (Fig. 4A). Further analysis of gene synteny within the *tapA* operon revealed two distinct sub-classes based on the presence or absence of *tapA*, as has been previously been noted for *B. cereus*, which contains two TasA paralogues but lacks *tapA* (Caro-Astorga *et al*., 2015). Highlighted on the phylogenetic tree are *B. amyloliquefaciens* TasA, *B. licheniformis* TasA and TasA and CalY from *B. cereus* that were chosen for further analysis (Fig. 4A). Each of these proteins are predicted to encode an N-terminal signal sequence and were used to establish: 1) whether orthologous TasA fibres assembled *in vitro* after purification of the predicted mature protein; and 2) if any fibres formed could cross-complement the *B. subtilis* Δ*tasA sinR* deletion strain.

**Figure 4:**
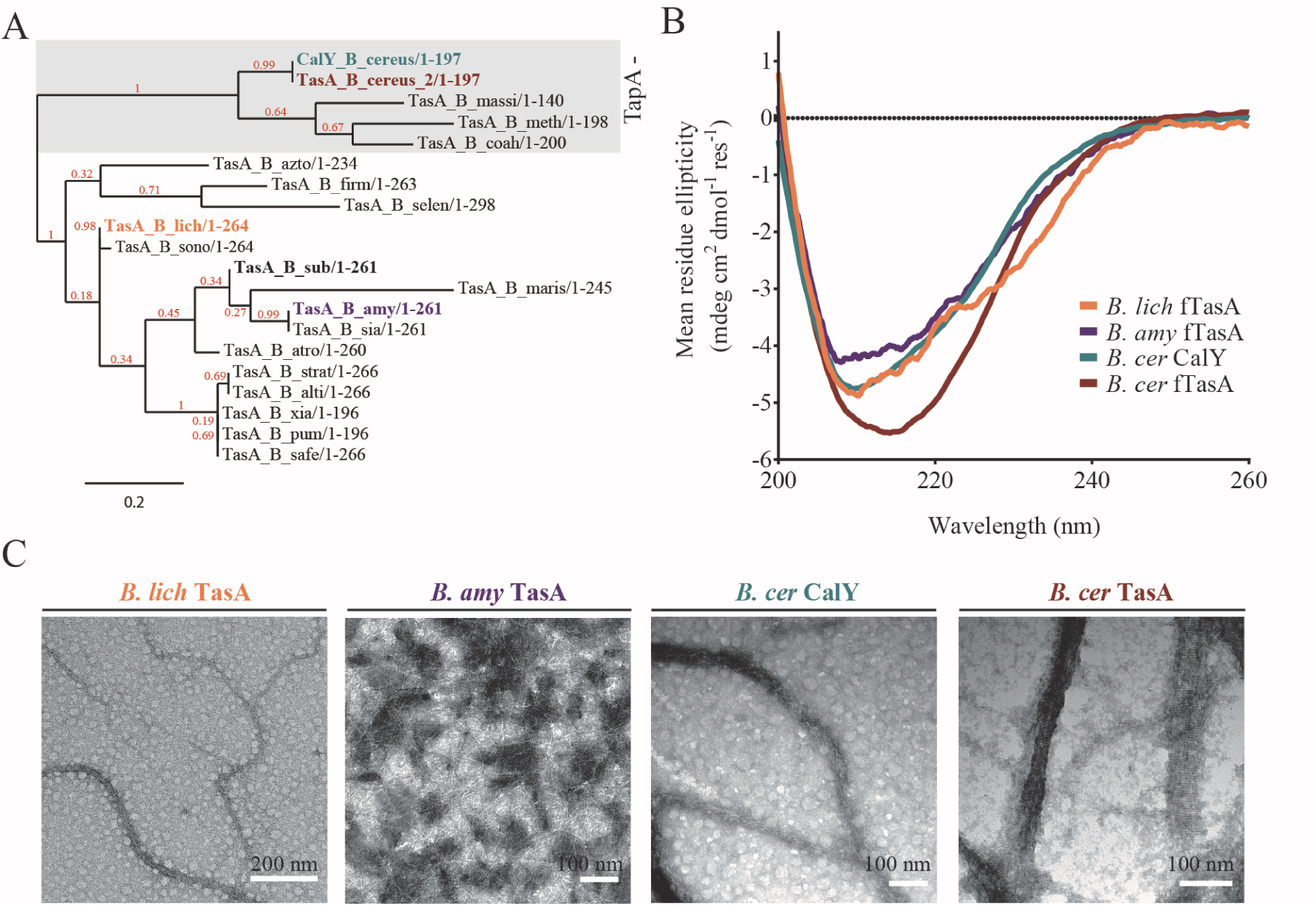
Characterisation of recombinant orthologous TasA proteins. (**A**) The phylogenetic tree was rooted using the midpoint method with the bootstrap value (red) given as a value between 0 and 1, where 1 is a high score. Highlighted are species chosen for subsequent analysis: *B. licheniformis* (orange), *B. amyloliquefaciens* (purple), *B. cereus* TasA (red) and CalY (green). For the protein sequence alignment and abbreviations of species names alongside accession numbers see Fig. S5. (**B**) Solution state circular dichroism spectra of recombinant *B. licheniformis, B. amyloliquefaciens, B. cereus* TasA and CalY. (**C**) Transmission electron microscopy images of recombinant orthologue TasA stained with uranyl acetate show the presence of fibres that are several micron in length and vary in width from 15 nm (*B. licheniformis*) to 25 nm (*B. amyloliquefaciens* and *B. cereus* CalY) and 60 nm (B. cereus TasA). A repeating unit at 4-5 nm is seen for all proteins.

The quality and identity of the recombinant TasA orthologous proteins was confirmed by SDS-PAGE (Fig. S6A) and mass spectrometry (Fig. S6B). CD spectroscopy indicated that the secondary structures of *B. licheniformis* and *B. amyloliquefaciens* TasA, and *B. cereus* CalY, were broadly similar to that of *B. subtilis* TasA, with a primary minimum at 208 nm, a shoulder at ~222 nm, and a maximum below 200 nm (Fig. 4B). In contrast, *B. cereus* TasA has a single broad minimum centred on ~216 nm (Fig. 4B), suggesting this protein may contain increased β-sheet content, although both the breadth and the intensity of the minimum suggest significant remaining contribution from helical elements. TEM imaging revealed that all of the orthologous proteins spontaneously self-assembled into fibres (Fig. 4C, S6C), and all showed evidence of a regular subunit repeat along the fibre axis of approximately 4-5 nm. The finding that proteins corresponding to the mature region of *B. cereus* TasA and CalY form fibres *in vitro* is consistent with previous data, which revealed the presence *in vivo* of extracellular fibres dependent on *tasA* and *calY* (Caro-Astorga *et al*., 2015) and with our findings that TapA is dispensable for TasA fibre formation *in vitro* (Fig. 3A). Through TEM imaging, we observed qualitative differences between the ability of the different proteins to form fibre bundles, with the *B. cereus* proteins TasA and CalY forming thick fibre bundles, *B. licheniformis* and *B. subtilis* TasA forming intermediate-diameter fibre bundles, and *B. amyloliquefaciens* forming a distributed mesh of thin fibres.

### Functionality of orthologous protein fibres in *B. subtilis*

To test the ability of the orthologous proteins to function in place of *B. subtilis* TasA fibres, 10 μg of each recombinant fibrous protein was exogenously added to the Δ*tasA sinR* mutant. The cells were then incubated under biofilm formation conditions. We determined that rugosity of the biofilm community could be recovered when the more closely related *B. amyloliquefaciens* and *B. licheniformis* TasA proteins were provided but not when either of the more divergent *B. cereus* proteins were supplied (Fig. 5A, S7A). This is in contrast to previously published data where expression of both *B. cereus calY* and *tasA*, alongside the signal peptidase *sipW*, was reported to recover biofilm formation to a *B. subtilis tasA* mutant (Caro-Astorga *et al*., 2015). Analysis of the stability of the protein fibres after incubation with spent cell-free culture supernatant revealed that *B. cereus* TasA fibres, like the *B. amyloliquefaciens* and *B. licheniformis* TasA fibres, were resistant to exoprotease degradation, while CalY fibres were susceptible (Fig. S7B-C). From our analyses of protein function we can conclude that either the interaction of TasA fibres with the *B. subtilis* matrix is dependent on the exact identity of the TasA fibres, suggesting specific molecular interactions with other matrix molecules, or that the subtle differences in the physiochemical properties of the TasA fibres may be influential in establishing rugosity in the bacterial biofilm. For example, after serially diluting the recombinant protein, and therefore shearing of the samples, recombinant *B. cereus* TasA was significantly less viscous than the equivalent samples of *B. licheniformis* and *B. subtilis* TasA (Fig. S7D-F) and we speculate that shearing of the samples breaks the thicker bundles observed in TEM. However, after allowing all samples to recover for 3 days, both *B. cereus* TasA and *B. licheniformis* TasA formed a gel at a lower concentration of protein than *B. subtilis* fTasA (Fig. S7D-F). This variability in the properties of the gels formed by the fibrous TasA orthologues may have implications for the mechanical properties of *in vivo* biofilms and the ability of one orthologue of TasA to substitute for another.

**Figure 5.**
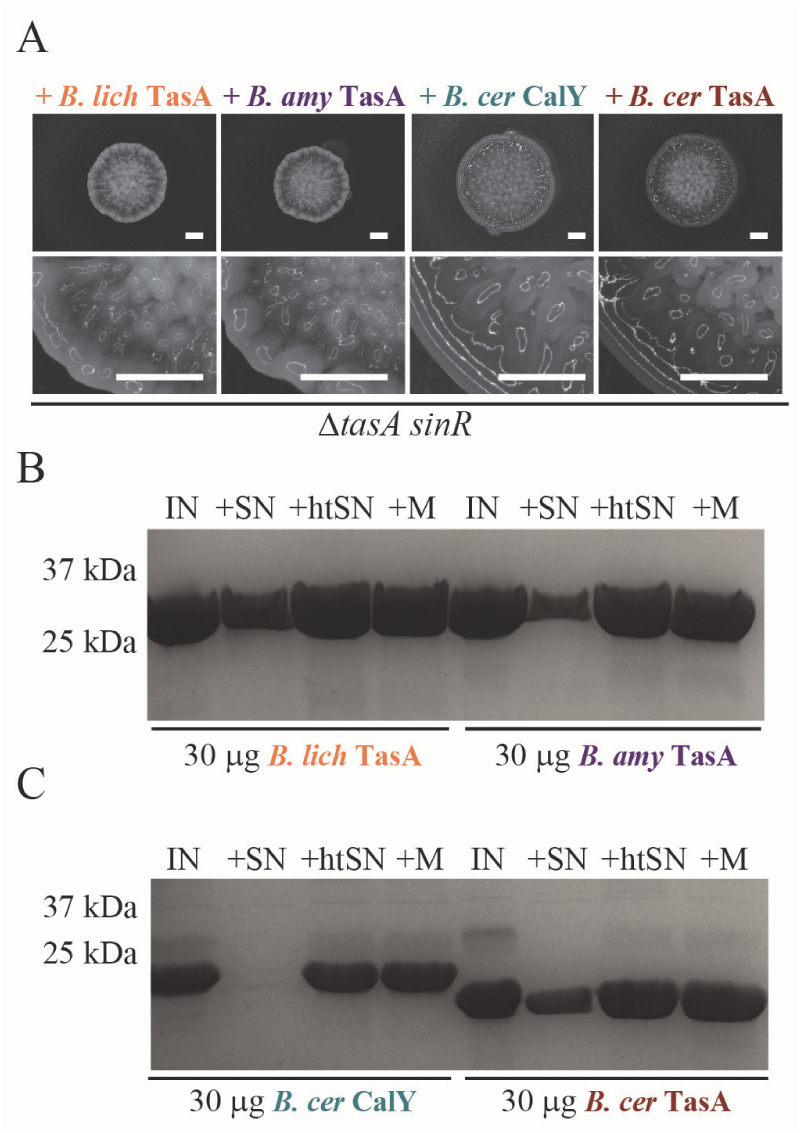
Complementation of *tasA sinR* null biofilm by recombinant orthologous TasA. (**A**) Biofilm phenotypes shown for Δ*tasA sinR* (NRS5248) after co-culture with 10 μg recombinant *B. licheniformis* TasA, *B. amyloliquefaciens* TasA and *B. cereus* TasA and CalY fibres. (**B-C**) Integrity of 30 μg *B. licheniformis, B. amyloliquefaciens* and *B. cereus* TasA and CalY proteins incubated for 24 hrs at 37°C analysed by SDS-PAGE. The protein (IN) was incubated with filtered spent supernatant collected from NCIB3610 (+SN) and supernatant subjected to heat inactivation at 100°C (+htSN) alongside media only control (+M).

## Discussion

We have demonstrated that recombinant fibrous TasA can return rugosity to a *B. subtilis* Δ*tasA sinR* deletion strain and shares the biological functionality of native TasA purified from *B. subtilis*. Biophysical analysis indicates that these fibres are assembled as a helical arrangement of globular units that lack the characteristic “cross-β” architecture of canonical amyloid-like fibres. The CD spectrum of the recombinant protein resembles that published previously for native TasA isolated directly from *B. subtilis* (Romero *et al*., 2010; Chai *et al*., 2013) and is suggestive of a predominantly helical secondary structure. Moreover, we have demonstrated that recombinant TasA can be rendered monomeric by the addition of a single amino acid to the N-terminus, and that this monomeric protein shares the same secondary structure as the fibrous form. This strongly suggests that the fibres comprise a linear assembly of these monomeric units, with no large structural rearrangement, although domain-swapping between monomers cannot be ruled out. Indeed, a repeating unit is visible along the length of the fibre axis, most clearly in the TEM images of recombinant fibres of the orthologous TasA protein from *B. cereus* where the protein subunits appear horizontally aligned across a fibre bundle, but also visible in all forms of TasA we have examined. Such a structure is not consistent with current structural models of amyloid-like fibrils, which comprise a single continuous hydrogen-bonded array along the long axis of the fibril. Taken together, our data indicate that TasA is unlikely to fall into the class of functional amyloid-like fibres.

We further found that our recombinant forms of TasA did not bind either Congo Red or ThT dyes that are commonly used to assess the formation of amyloid-like fibres. Moreover, our protein extracts from *B. subtilis* showed dye binding activity irrespective of whether TasA was present or not. Caution should be taken when inferring the formation of amyloid-like fibres from enhanced fluorescence in the presence of ThT, which also exhibits enhanced fluorescence in the presence of globular proteins such as bovine serum albumin (Freire *et al*., 2014), human serum albumin (Sen *et al*., 2009) and acetylcholinesterase (De Ferrari *et al*., 2001); in the presence of amorphous aggregates of lysozyme and bovine serum albumin (Yang *et al*., 2015), and amorphous aggregates formed by a thrombin-derived C-terminal peptide (Petrlova *et al*., 2017); and in the presence of non-amyloid wormlike aggregates of an artificial dimer of an Aβ peptide (Yamaguchi *et al*., 2010). Conversely, ThT does not exhibit enhanced fluorescence in the presence of, for example, cross-β fibrils formed by poly-L-lysine (Benditt, 1986; LeVine, 1999). Congo Red is similarly promiscuous (Howie and Brewer, 2009), although the observation of green birefringence under cross-polarisers is one of the identifying characteristics of amyloid deposits *in vivo*. Thus, Congo Red binding and enhanced ThT fluorescence should be considered only suggestive, but not indicative, of amyloid-like fibre formation.

The widespread nature of functional amyloid fibres in bacterial biofilms has been hypothesized, and a well-characterised example is the curli fibres of *E. coli, Enterobacter cloacae*, and *Salmonella* spp (Evans and Chapman, 2014). These show a CD spectrum, dye-binding behaviour, enhanced stability and proteolytic insensitivity that are consistent with an amyloid-like β-sheet structure, but solid-state NMR data suggests an architecture comprising stacked β-helical subunits (Shewmaker *et al*., 2009), a structural motif commonly employed by bacteria (Kajava and Steven, 2006). Many amyloid-like fibres formed *in vitro* from proteins associated with disease show an in-register parallel cross-β arrangement (Margittai and Langen, 2008); recently however native Tau filaments extracted from the brain of an Alzheimer’s Disease patient have been demonstrated to form an elaborate mixed β-helix/cross-β structure formed of in-register, parallel β-strands (Fitzpatrick *et al*., 2017). Thus, both cross-β and β-helix architectures may be characteristic of amyloid fibres, and curli fibres may still be considered as “amyloid-like”.

Making the correct distinction between amyloid-like and non-amyloid fibrous proteins is more than a semantic argument: a number of papers have drawn a link between functional amyloid-like fibres formed by bacteria and their relevance to human disease (Epstein and Chapman, 2008; Chai *et al*., 2013; Evans and Chapman, 2014), for example, in the determination of the mechanistic details of self-assembly, or in the possible discovery of new therapeutics. As the amyloid-like fibre macrostructure is thought to be a ‘generic’ property deriving from the chemical structure of the polypeptide backbone that is common to all proteins and peptides – and thus to a large extent independent of primary sequence, although this will influence overall fibre morphology - small drug molecules that target the generic amyloid fold may have widespread applicability in a number of devastating human diseases. Thus it is important to make the distinction between non-amyloid fibrous assemblies and amyloid-like fibres appropriately.

The fibrous nature of TasA likely imparts mechanical rigidity to the biofilm, thereby restoring the highly wrinkled architecture characteristic of the *ΔtasA sinR* deletion strain. As indicated above it is unclear why neither fTasA nor nTasA(+) can recover biofilm architecture to the single *tasA* deletion and furthermore, why expression of a *sipW-tasA* construct is required for genetic complementation. Since SinR has pleiotropic roles in biofilm formation (Vlamakis *et al*., 2013; Cairns *et al*., 2014) it may be that overproduction of the biofilm polysaccharide compensates for the loss of native regulation that intricately controls native TasA production in space and time (Vlamakis *et al*., 2008). Our results also indicate that when in a fibrous form, TasA does not require the TapA protein to fulfil its function, which contradicts previous reports suggesting that TapA is an accessory protein required for correct TasA assembly and localisation (Romero *et al*., 2011). Therefore the role played by TapA in biofilm formation, while evidently essential (Chu *et al*., 2006), is unclear. It may be that while TapA is not essential for TasA fibre formation *in vitro*, it functions as a chaperone *in vivo* to aid the transition of monomeric TasA into a fibrous state. This hypothesis is consistent with the overall reduction in the level of TasA and the corresponding reduction in the number of TasA fibres observed in the *tapA* mutant (Romero *et al*., 2011). Moreover it is consistent with the demonstration that monomeric TasA, but not fibrous TasA, is susceptible to degradation by the extracellular proteases.

A non-amyloid-like structure for TasA is possibly beneficial in the context of the *B. subtilis* biofilm; amyloid-like self-assembled fibres are very stable, with curli fibres, for example, requiring treatment with concentrated acid solutions to drive disassembly (Chapman *et al*., 2002). Curli fibres also appear to form a brittle matrix which, once fractured, does not recover (Serra *et al*., 2013). In contrast, we have shown that the gelation properties of fibrous TasA solutions recover after shear (Fig. SI 5D-F), suggesting that *in vivo* the biofilm matrix could be remodelled in response to mechanical environmental perturbations. The TasA fibres may also be in equilibrium with the monomeric form of the protein, which would allow dynamic restructuring of the biofilm in response to environmental changes. As the fibrous form of the protein confers protection against degradation by extracellular proteases whereas the monomeric protein is degraded, an appropriate secretion of monomeric protein and/or proteases could provide dynamic control of biofilm elasticity and structure.

## Materials and Methods

### Growth conditions

*E. coli* and *B. subtilis* were routinely grown in Lysogeny Broth (LB) media (10 g NaCl, 5 g yeast extract and 10 g tryptone per litre) or plates supplemented with the addition of 1.5% (w/v) select agar (Invitrogen). Samples were grown at 37°C unless stated otherwise. When required, antibiotics were used at the following concentrations: ampicillin (100 μg ml^−1^), spectinomycin (100 μg ml^−1^) and chloramphenicol (5 μg ml^−1^). For biofilm assays MSgg minimal media was used (5 mM KH_2_PO_4_ and 100 mM MOPS at pH 7 supplemented with 2 mM MgCl_2_, 700 μM CaCl_2_, 50 μM MnCl_2_, 50 μM FeCl_3_, 1 μM ZnCl_2_, 2 μM thiamine, 0.5% glycerol, 0.5% glutamate). When appropriate isopropyl β-D-1-thiogalactopyranoside (IPTG) was added at the indicated concentration. For protein production auto-induction media (6 g Na_2_HPO_4_, 3 g KH_2_PO_4_, 20 g Tryptone, 5 g yeast extract, 5 g NaCl, 10 ml 60% v/v glycerol, 5 ml 10% w/v glucose and 25 ml 8% w/v lactose per litre at a 1:1000 volume ratio (supplemented with 100 μg/ml ampicillin)) was used (Studier, 2005).

### Strain construction

A complete list of *E. coli* and *B. subtilis* strains used in this study can be found in Table S1. Plasmids and primers are detailed in Tables S2 and S3 respectively. All *B. subtilis* strains used for physiological assays were derived from the wild-type laboratory isolate NCIB3610 and constructed using standard protocols. SSP1 phage transductions for DNA transfer into *B. subtilis* NCIB3610 were carried out as previously described (Verhamme *et al*., 2007).

### Plasmid construction and mutagenesis

Construction of an in-frame *tasA* deletion in NCIB3610 was achieved using the pMiniMAD (Patrick and Kearns, 2008) temperature sensitive allelic replacement vector pNW1448 (pMiniMAD-Δ*tasA*). The plasmid was constructed by PCR amplification of two fragments: the 511 bp upstream of *tasA* including the first 6 bp of *tasA* coding sequence and a second fragment covering the last 3bp of the *tasA* coding sequence, the stop codon and the 512 bp downstream using primer pairs NSW2005/NSW2006 and NSW2007/NSW2008 respectively. The PCR fragments were each digested with SalI/EcoRI and simultaneously ligated into the pMiniMAD plasmid that was digested with the same restriction sites to yield plasmid pNW1448. Plasmid pNW1448 was introduced into 168 and then transferred to NCIB3610 using phage transduction. The *tasA* deletion was introduced into the *B. subtilis* chromosome using the method described previously (Arnaud *et al*., 2004). After homologous recombination and selection for loss of the pMiniMad plasmid, two morphologically distinct isolates carrying the desired deletion in *tasA* were identified. Whole genome sequencing (see below) was used to genotype the isolates in an unbiased manner. Analysis of single nucleotide polymorphisms (Table S4) revealed one strain carried a short duplication of the *sinR* coding region effectively yielding a Δ*tasA sinR* double mutant (NRS5248) while the other was a single *ΔtasA* strain (NRS5267).

The *tapA* in frame deletion was generated via the pMAD protocol as above, with amplification of the 395 bp upstream fragment using primers NSW1308 and NSW1332 and 641 bp downstream fragment using primers NSW1333 and NSW1334. The two PCR fragments were digested BamHI/SalI and EcoRI/SalI respectively and ligated into the intermediate plasmid pUC19 yielding pNW686, and was subsequently moved into pMAD to generate pNW685. Plasmid pNW685 was introduced into *B. subtilis* 168, generating strain NRS3789, and transferred to NCIB3610 using phage transduction. The same 168 strain was used to generate the Δ*tapA* Δ*tasA* and Δ*tapA* Δ*tasA sinR* strains by transferring via phage to Δ*tasA* (NRS5267) and Δ*tasA sinR* (NRS5248) respectively.

Genetic complementation of Δ*tasA* and Δ*tasA sinR* was achieved by PCR amplification of the *tasA* (using primers NSW1857 and NSW1858) and *sipW-tasA* (using primers NSW2218 and 2219) regions from NCIB3610. Both were cut using SalI/SphI restriction enzymes and ligated into the pDR183 (pNW1434) and pDR110 (pNW1432 and pNW1619) vector. Plasmid pNW1434 pDR183 was introduced to 168 and transferred to Δ*tasA sinR* (NRS5255). Plasmids pNW1432 and pNW1619 were introduced into 168 and transferred to Δ*tasA* (NRS5276) and Δ*tasA sinR* (NRS5248) using phage transduction.

Genetic complementation of the triple Δ*tapA* Δ*tasA sinR* (NRS5749) mutant was performed using the whole *tapA-sipW-tasA* operon which was amplified from NCIB3610 using primers NSW1896 and NSW2219, cut SalI/SphI and ligated into pDR110 to generate pNW1804 which was introduced by phage transduction via 168 at the ectopic *amyE* location on the chromosome.

Protein purification was achieved using GST fusion constructs. The *tasA* overexpression plasmid pNW1437 (pGex-6-P-1-TEV-*tasA*_(28-261)_) was generated by amplifying the *tasA*_(28-261)_ coding region from *B. subtilis* NCIB3610 genomic DNA using primers NSW660 and NSW661 and insertion into the vector pGEX-6P-1 cleaved BamHI/XhoI yielding the vector pNW543. The TEV protease cleavage site was next introduced by site-directed mutagenesis using primers NSW1892 and NSW1893 to give pNW1437. Amino acids were introduced at the N-terminal end of *tasA* also by site-directed mutagenesis; primer pairs are indicated in Table S2. The constructs used to purify the TasA orthologue proteins were generated in a similar manner from genomic DNA isolated from *B. cereus* ATCC14579, *B. licheniformis* ATCC14580 and *B. amyloliquefaciens* FZB42 and likewise primers used for amplification are detailed. The plasmids were used to transform BL21 (DE3) *E. coli* strain for protein production.

### Genome sequencing

Whole genome sequencing and bioinformatics analysis of strains NCIB3610, NRS5248 and NRS5267 was conducted by MicrobesNG (http://microbesng.uk) which is supported by the BBSRC (grant number BB/L024209/1). Three beads were washed with extraction buffer containing lysozyme and RNase A, incubated for 25 min at 37°C. Proteinase K and RNaseA were added and incubated for 5 minutes at 65°C. Genomic DNA was purified using an equal volume of SPRI beads and resuspended in EB buffer. DNA was quantified in triplicates with the Quantit dsDNA HS assay in an Ependorff AF2200 plate reader. Genomic DNA libraries were prepared using Nextera XT Library Prep Kit (Illumina™, San Diego, USA) following the manufacturer’s protocol with the following modifications: 2 ng of DNA instead of one were used as input, and PCR elongation time was increased to 1 min from 30 seconds. DNA quantification and library preparation were carried out on a Hamilton Microlab STAR automated liquid handling system. Pooled libraries were quantified using the Kapa Biosystems Library Quantification Kit for Illumina on a Roche light cycler 96 qPCR machine. Libraries were sequenced on the Illumina HiSeq using a 250 bp paired end protocol. Reads were adapter trimmed using Trimmomatic 0.30 with a sliding window quality cut-off of Q15 (Bolger *et al*., 2014). De novo assembly was performed on samples using SPAdes version 3.7 (Bankevich *et al*., 2012) and contigs were ordered using Abacas (Assefa *et al*., 2009) and annotated using Prokka 1.11 (Seemann, 2014). Reads were aligned to the reference 168 genome (accession number: NZ_CM000487.1) using BWA-Mem 0.7.5 and processed using SAMtools 1.2 (Li *et al*., 2009). Variants were called using VarScan 2.3.9 with two thresholds, sensitive and specific, where the variant allele frequency is greater than 90% and 10% respectively. The effects of the variants were predicted and annotated using SnpEff 4.2 (Koboldt *et al*., 2009) (Table S4).

### Protein production and purification

The pGEX-6-P-1 GST-gene fusion system was used in the *E. coli* BL21 (DE3) strain for protein production (GE Healthcare™). After the required plasmid was introduced into BL21(DE3), a 5 ml LB culture (supplemented with 100 μg/ml ampicillin) was grown overnight at 37°C and used to inoculate 1 L of auto-induction media at 1/1000 dilution. The cultures were incubated at 37°C with 130 rpm shaking until optical density at 600 nm was approximately 0.9 at which point the temperature was lowered to 18°C and cultures were grown overnight. Cells were harvested by centrifugation at 4000 g for 45 minutes and the cell pellet was suspended in 25 ml of purification buffer (25 mM Tris-HCl, 250 mM NaCl, pH 7.5) supplemented with Complete EDTA-free Protease inhibitor (Roche) then lysed using an Emulsiflex cell disrupter (Avestin™) with 3 passes made at ~15000 psi or sonication at 25% for 6 minutes. Cell debris was removed by centrifugation at 27000 g for 35 minutes. The supernatant was removed and added to 450 μL of Glutathione Sepharose 4B beads (GE Healthcare™) and incubated on a roller for 3 hours at 4°C. The protein-bead mix was loaded onto disposable gravity flow columns (Bio-Rad™) and washed 3 times with 25ml of purification buffer. Beads were collected from the column and suspended in 25 ml of purification buffer supplemented with 1 mM DTT and 0.5 mg TEV protease and incubated on roller at 4°C overnight. The flow-through was then added to 300 μL GST beads and 250 μL Ni-NTA beads (Qiagen™) and incubated for 2 hours at 4°C. Final pass through column removes beads and flow-through is concentrated using 10kDa Vivaspin™. For biophysical experiments performed at Edinburgh University, buffer exchange into 25 mM phosphate buffer (pH 7) was performed using same concentrators. Purity was determined by SDS-PAGE and molecular mass determined by loading 80 μg onto a qTOF liquid chromatography mass spectrometry performed by the FingerPrints Proteomics Facility at the University of Dundee.

### Native extraction from *B. subtilis*

Method adapted from (Romero *et al*., 2010). Briefly, cells from the *eps sinR* double mutant and *eps sinR tasA* triple mutant were grown in 1L Msgg at 37° at 130 rpm for 20 hours from an OD_600_ of 0.02. Cells were pelleted at 5,000 g for 30 minutes and the media discarded. Cells were centrifuged twice with 25 ml extraction buffer (5mM KH_2_PO_4_, 2 mM MgCl_2_, 100mM MOPS (pH 7), 1M NaCl with Roche Protease Inhibitor cocktail) and the supernatant filtered through a 0.4 μm filter. Ammonium sulphate was added to make 30% in final volume and incubated with stirring at 4°C for 1 hour. The supernatant was then centrifuged at 20,000 g for 10 minutes to remove precipitated proteins and dialysed twice in 5 L 25 mM phosphate buffer (pH 7) at room temperature for 1 hour each and then 4°C overnight.

### Transthyretin fibre preparation

TTR fibres were prepared as previously described (Schor *et al*., 2015). Briefly, 0.8 mg of the peptide was dissolved in 200 μL 25mM phosphate buffer (pH 7) for *ex vivo* complementation and 20% (w/v) acetonitrile (pH 5) for X-ray diffraction pattern collection. Sample was sonicated for 10 minutes and combined with 10 μL of TTR seeds and incubated at 60°C for 5 hours.

### Biofilm phenotypes, ex vivo complementation and protein collection

To characterise biofilm phenotype samples were set up as detailed previously (Branda *et al*., 2001) briefly, 10 μL of LB culture grown to mid-exponential phase was spotted onto solidified MSgg media and incubated for 2 days at 30°C. The resultant colonies were imaged using a Leica MZ16 stereoscope. For *ex vivo* complementation, 10 μg of recombinant protein, 30 μg native extract or 10 μL TTR where indicated was pipetted with cells immediately prior to spotting. To release all biofilm proteins for subsequent analysis, the biofilm was resuspended in 500 μL BugBuster Master Mix (Novagen) followed by sonication and agitation for 20 mins at room temperature. Insoluble debris was removed by centrifugation at 17,000 g for 10 mins at 4°C.

### SDS-PAGE

SDS-polyacrylamide gel electrophoresis (PAGE) was performed using 10 μg of purified TasA protein and 4X loading buffer (6.2 g SDS, 40 ml 0.5 M Tris pH 6.8, 6.4 ml 0.1 M EDTA, 32 ml 100% glycerol, 1 mg Bromophenol blue). Samples were heated at 99°C for 5 minutes prior to loading on the gel and were run on a standard 14 % polyacrylamide SDS-PAGE at 200 V for 60 min, before staining with InstantBlue (Expedeon™).

### Immunoblot Analysis

Samples were separated by SDS-PAGE and transferred onto a PVDF membrane (Millipore™) by electroblotting at 100 mA for 75 minutes. The membranes were blocked with 3% (w/v) milk in 1xTBS overnight at 4°C with shaking followed by 1 hr incubation with primary antibody (TasA (1:25000 v/v) as indicated) diluted in 3% (w/v) milk in 1x TBS. This was followed by 3 washes of 10 minutes each with 1x TBS and 2% (v/v) Tween20 and subsequent 45 minute incubation with goat anti-rabbit conjugated secondary antibody (1:5000 v/v) (Pierce™). Membrane was washed 3 times for 10 mins with TBST then developed by ECL incubation and exposing to X-ray film (Konica™) using the Medical Film Processor SRX-101A (Konica™). This is with the exception of the data shown in Figure 4C which was developed as detailed above and visualised using GeneGnomeXRQ (Synegene™).

### Size-Exclusion Chromatography

Monomeric TasA was examined by size-exclusion chromatography using either Superdex 5/150 or 10/300 GL increase column as indicated (GE Healthcare) on an ÄKTA FPLC system using 25 mM Tris-HCl, 250 mM NaCl, pH 7 buffer. Column was calibrated using conalbumin (75000 Da), ovalbumin (44000 Da), carbonic anhydrase (29000 Da), ribonuclease A (13700 Da) and aprotinin (6500 Da) and void volume was calculated using blue dextran 2000 (GE Healthcare).

### Exoprotease stability

PY79, PY79 Δ6 and PY79 Δ7 and/or NCIB3610 were grown to an OD_600_ of ~2.5 in 25 ml MSgg growth media at 37°C with 130 RPM shaking overnight. The cultures were normalised to same OD_600_ and 5 ml was centrifugation at 3750 *g* for 15 mins at 4 °C to pellet cells. The culture supernatant was collected and filtered through a 0.22 μM filter (Milipore) to remove residual cells. Aliquots of the culture supernatant generated by NCIB3610 and PY79 were heated inactivated at 100°C for 10 minutes as requried. 15 μL of each cell-free culture supernatant was incubated with 30 μg recombinant protein alongside a media only-control at 37°C for 24 hours. The integrity of the protein was analysed by SDS-PAGE alongside a non-incubated sample of recombinant protein as a loading control.

### Protein Sequence Alignment

TasA orthologues were identified by BLASTP (Altschul *et al*., 1990; Altschul *et al*., 1997) using the protein sequence of TasA from *B. subtilis* as the query. TasA protein sequences were aligned using Clustal Omega with the default settings (Sievers *et al*., 2011). The aligned sequences were imported and manually coloured for homology as indicated in the legend in Microsoft Word. The signal sequences were predicted using the SignalP v4.1 server and are indicated by underline (Petersen *et al*., 2011). A maximum likelihood tree was calculated from the Clustal Omega alignment using the phylogeny.fr platform (Dereeper *et al*., 2008), Gblocks was used to eliminate divergent and poorly aligned segments for tree construction (Castresana, 2000). The tree was estimated using the PhyML algorithm (Guindon and Gascuel, 2003) with mid-point rooting, using a WAG substitution model (Whelan and Goldman, 2001) and bootstrapping procedure set to 100 replicates. The outputted tree was visualised using TreeDyn (Chevenet *et al*., 2006).

### Protein Precipitation of TasA for Mass spectrometry

A strain carrying an IPTG inducible copy of the *tasA* gene (NRS5313) in a Δ*tasA* background was grown in 200 ml Msgg at 37°C until OD_600_ of 1 in the presence of 1 mM IPTG. The culture supernatant was collected and separated from cell fraction by centrifugation at 5000 *g* for 30 minutes at 4°C with iterative removal of the supernatant into a fresh tube for 4 rounds. 40 ml of the clarified supernatant was precipitated overnight with 6.25% (w/v) trichloroacetic acid (Sigma™) at 4°C and the precipitated proteins were recovered by centrifugation as before. The protein pellet was washed 5 times with 1 ml ice-cold dH_2_0 and air dried for 1 hour (protocol modified from Cianfanelli *et al*. 2016). The protein pellet was suspended in 50 μL 2x laemmli buffer and separated by SDS-PAGE on a 14% gel alongside *in vitro* purified fTasA protein as a size marker. The section of the lane at the position expected to contain mature TasA was excised and analysed by mass spectrometry.

### Mass Spectrometry

Samples were processed prior to overnight (16 h) trypsin digestion (Modified Sequencing Grade, Pierce). Peptides extracted from gel and dried in SpeedVac (Thermo Scientific™). Peptides re-suspended 50 μl 1% formic acid, centrifuged and transferred to HPLC vial. 15 μl of this was typically analysed on the system. The peptides from each fraction were separated using a mix of buffer A (0.1% formic acid in MS grade water) and B (0.08% formic acid in 80% MS grade CH_3_CN). The peptides from each fraction were eluted from the column using a flow rate of 300 nl/min and a linear gradient from 5% to 40% buffer B over 68 min. The column temperature was set at 50 °C. The Q Exactive HF Hybrid Quadrupole-Orbitrap Mass Spectrometer was operated in data dependent mode with a single MS survey scan from 335-1,800 m/z followed by 20 sequential *m/z* dependent MS2 scans. The 20 most intense precursor ions were sequentially fragmented by higher energy collision dissociation (HCD). The MS1 isolation window was set to 2.0 Da and the resolution set at 60,000. MS2 resolution was set at 15,000. The AGC targets for MS1 and MS2 were set at 3e^6^ ions and 5e^5^ ions, respectively. The normalized collision energy was set at 27%. The maximum ion injection times for MS1 and MS2 were set at 50 ms and 100 ms, respectively. Exactive HF Hybrid Quadropole .RAW data files were extracted and converted to mascot generic files (.mgf) using MSC Convert. Extracted data then searched against the Local peptide database containing the relevant TasA sequence using the Mascot Search Engine (Mascot Daemon Version 2.3.2).

### Fibre formation and X-ray Diffraction

To prepare samples for X-ray diffraction, 5 μL of recombinant at ~ 5 mg/ml fTasA was suspended between 2 borosilicate glass capillaries (Harvard Apparatus) and allowed to dry (Makin & Serpell, 2005). The dried fibres were mounted onto a Rigaku M007HF X-ray generator equipped with a Saturn 944HG+ CCD detector, and images collected with 60s exposures at room temperature. Diffraction patterns were inspected using Ipmosflm CCP4} and then converted to TIFF format. CLEARER (Sumner Makin *et al*., 2007) was used to measure the diffraction signal positions.

### Circular Dichroism (CD) Spectroscopy

All CD measurements were performed using a Jasco J-810 spectropolarimeter. Solution-state samples were measured at a protein concentration of 0.2 mg/ml (in 25 mM phosphate buffer) in a 0.1 cm quartz cuvette. A scan rate of 50 nm/s was used, with a data pitch of 0.1 nm and a digital integration time of 1 s. Twenty scans were accumulated and averaged to produce the final curve.

### Transmission Electron Microscopy (TEM) imaging

A 5 μl droplet of 0.02 mg/ml protein solution was pipetted onto a carbon-coated copper grid (TAAB Laboratories Equipment Ltd) and left for 4 minutes before being wicked away from the side with filter paper. Subsequently, a 5 μl droplet of 2% (w/v) uranyl acetate was pipetted onto the grid and left for 3 minutes before being wicked away from the side with filter paper. The stained grids were imaged using a Philips/FEI CM120 BioTwin transmission electron microscope and ImageJ software was used for image analysis.

### Thioflavin T binding kinetics

Protein samples were diluted to 3 mg/ml in 25 mM phosphate buffer. 200 μL of protein was added into the wells of a Corning NBS 96-well plate (Corning 3641). ThT was added to a final concentration of 20 μM. The plates were sealed with a transparent film and put into a BMG Fluostar plate reader at 37°C as indicated. Measurements of ThT fluorescence were taken every 5 minutes for a period of 8 hours for mTasA and fTasA, the median of these values in represented in Fig S1G. For nTasA(+), nTasA(−) and controls, only a single read was taken.

### Congo Red Binding Assay

A stock solution of 2 mg/ml Congo Red (Sigma-Aldrich 75768) was prepared in phosphate buffer and filtered three times using a 0.22 μm syringe filter (Millipore). 2 mg/ml bovine insulin (Sigma-Aldrich I5500) was prepared in MilliQ water adjusted to pH 1.6 using concentrated HCL. The insulin sample was incubated overnight at 60°C. 60 μL of each protein sample was added to a cuvette containing 1 mL of buffer and 10 μL of the Congo Red stock solution. The samples were then allowed to incubate at room temperature for 30 minutes. A control spectrum containing only Congo Red was measured where 10 μL of the Congo Red stock solution was added to 1 mL of buffer plus an additional 60 μL of buffer (to match the amount of protein added to each cuvette). Since the nTasA(+/-) samples contained multiple components, a UV-vis spectrum (Cary 1E spectrophotometer) from 800 to 200 nm was measured and the relative absorbance peaks at 280 nm was used to ensure equal amounts of protein were measured between the two samples. The Congo Red spectra were acquired over a wavelength range of 400-700 nm.

### Mean square displacement via bead tracking

A 1 μL aliquot of carboxylate-modified polystyrene, fluorescent yellow-green latex beads with a mean particle size of 1 μm (Sigma-Aldrich, L4655) was diluted into 1 mL of phosphate buffer. 5 μL of this stock solution was added to the protein solution and gently mixed to disperse the particles. 80 μL of the bead and protein solution was placed on a cavity slide (Brand GmBH, 0.6-0.8 mm depth) and sealed with a coverslip using nail varnish. Movies of the motion of the particles were taken using a Nikon Eclipe Ti microscope equipped with a Hamamatsu Orca-Flash 4.0 CCD camera. Images were acquired using μ-manager software at a framerate of 10 fps (Edelstein *et al*., 2010). Movies were then analysed using TrackPy (available from github.com/soft-matter/trackpy).

## Acknowledgments

Work was supported by the Biotechnology and Biological Sciences Research Council [BB/L006804/1; BB/L006979/1; BB/M013774/1; BB/N022254/1]. EE and CE are supported by the Wellcome Institutional Strategic Support Fund (Award no. 097818/Z/11). We would like to acknowledge the FingerPrints Proteomics Facility and the X-ray Crystallography Facility at the University of Dundee (which is supported by Wellcome (Award no. 094090)). We are grateful to Ms. Ho, Dr. Hobley and Dr. Ostrowski for contributing plasmids and a strain used in this study.

---

**Table S1.** *B. subtilis* and *E. coli* strains used in this study

| Strain | Relevant genotype /Description | Source / Construction<sup>b,c</sup> |
| --- | --- | --- |
| MC1061 | *E. coli* F'*lacI*Q *lacZ*M15 *Tn10* (*tet*) | *E. coli* Genetic Stock Centre |
| BL21 | F<sup>−</sup> *ompT* *hsdSB*(rB<sup>−</sup>, mB<sup>−</sup>) *gal dcm* (DE3) | (Studier and Moffatt, 1986) |
| NCIB3610 | Prototrophic wild-type strain | BGSC |
| 168 | *trpC2* | BGSC |
| NRS5048 | 168 pMiniMAD-*ΔtasA* | pNW1448 → 168 |
| NRS5267 | NCIB3610 *ΔtasA* | SPP1 NRS5048 → 3610 |
| NRS5276 | NCIB3610 *ΔtasA lacA::P<sub>IP</sub>TG<sup>−</sup>-tasA* | pNW1434 → NRS5267 |
| NRS5313 | NCIB3610 *ΔtasA amyE::P<sub>IP</sub>TG<sup>−</sup>-sipW-tasA* | pNW1619 → NRS5267 |
| NRS5316 | NCIB3610 *ΔtasA amyE::P<sub>IP</sub>TG<sup>−</sup>-sipW<sub>TAA</sub>-tasA* | pNW1631 → NRS5267 |
| NRS5248 | NCIB3610 *ΔtasA sinR<sub>(Phe65_Iso71dup)</sub>* | SPP1 NRS5048 → 3610 |
| NRS5255 | NCIB3610 *ΔtasA sinR amyE::P<sub>IP</sub>TG<sup>−</sup>-tasA (spc)* | pNW1432 → NRS5248 |
| NRS1661 | 168 *epsA::pBL584Φ(P<sub>IP</sub>TG<sup>−</sup>-*epsA*) (cml)* | pBL584 → 168 |
| NRS5421 | NCIB3610 *ΔtasA sinR epsA::pBL584Φ(P<sub>IP</sub>TG<sup>−</sup>-*epsA*) (cml)* | SPP1 NRS1661 → NRS5248 |
| NRS3789 | 168 containing pNW685 | pNW685 → 168 |
| NRS5749 | NCIB3610 *ΔtapA ΔtasA sinR* | SPP1 NRS3789 → NRS5248 |
| NRS5760 | 168 *amyE::P<sub>IP</sub>TG<sup>−</sup>-tapA-sipW-tasA-lacI (spc)* | pNW1804 → 168 |
| NRS5763 | NCIB3610 *ΔtapA ΔtasA sinR amyE::P<sub>IP</sub>TG<sup>−</sup>-tapA-sipW-tasA-lacI (spc)* | SPP1 NRS5760 → NRS 5749 |
| NRS2450 | NCIB3610 *eps(A-O)::tet* | (Ostrowski *et al.*, 2011) |
| NRS1235 | 168 *sinR::cat* | NSW laboratory stocks |
| NRS5422 | *eps(A-O)::tet sinR::cat* | SPP1 NRS1235 → NRS2449 |
| NRS1858 | 168 *sinR::kan* | (Kiley and Stanley-Wall, |
| NRS5931 | *eps(A-O)::tet sinR::kan tasA::spc* | SPP1 1858 → NRS2450 |
| PY79 Δ7 | PY79 *nprE aprE epr mpr nprB vpr bpr* | BGSC KO7 (1A1133) |
| PY79 Δ6 | PY79 *nprE aprE epr mpr nprB vpr* | BGSC KO6 |

1. Drug resistance cassettes are indicated as follows: *spc*, spectinomycin, *amp*: ampicillin, *cml*: chloramphenicol, *kan*:: kanamycin 2. BSGC represents the *Bacillus* genetic stock centre. 3. The direction of strain construction is indicated with plasmid DNA or phage (SPP1) (→) recipient strain.

**Table S2.**
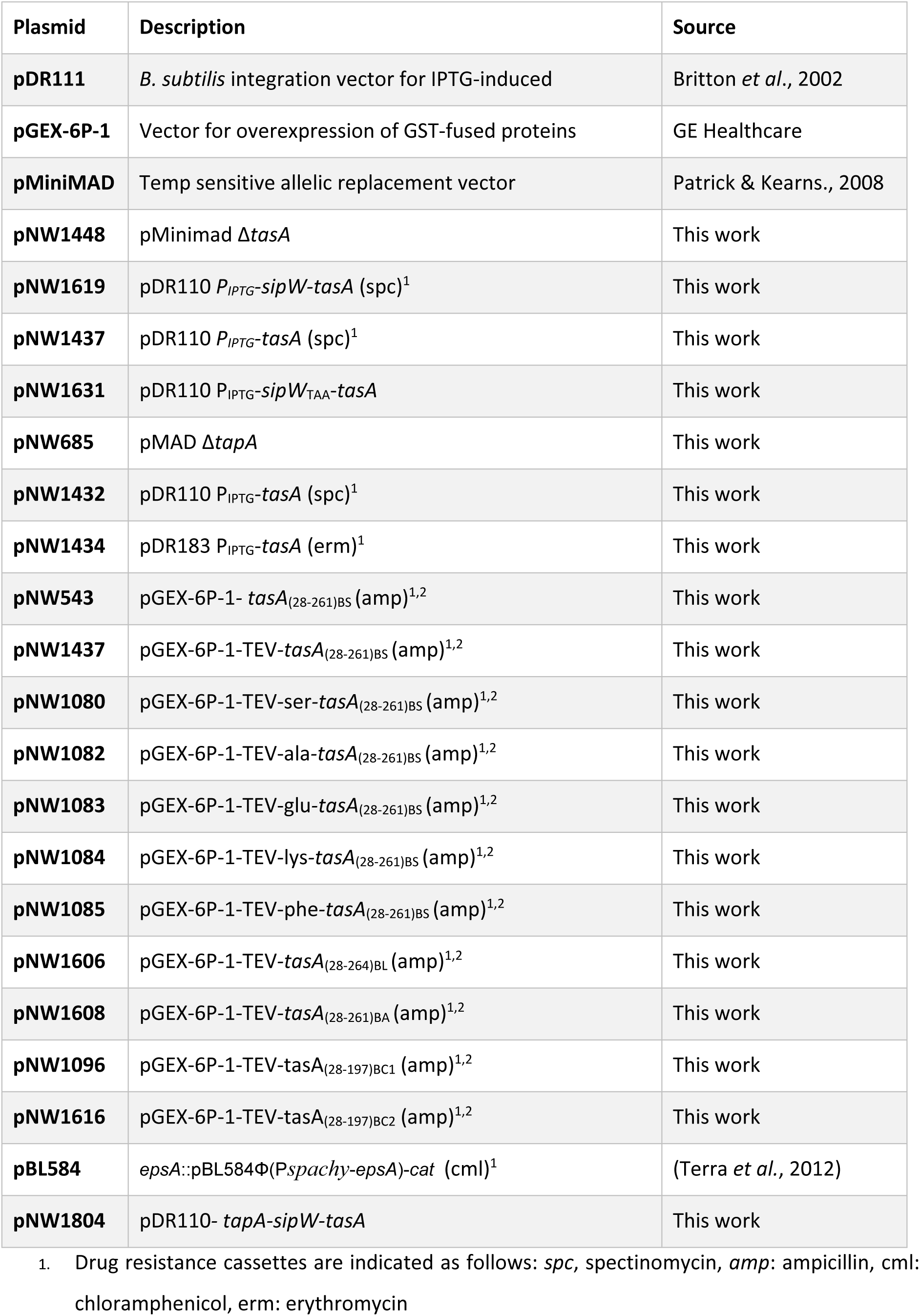
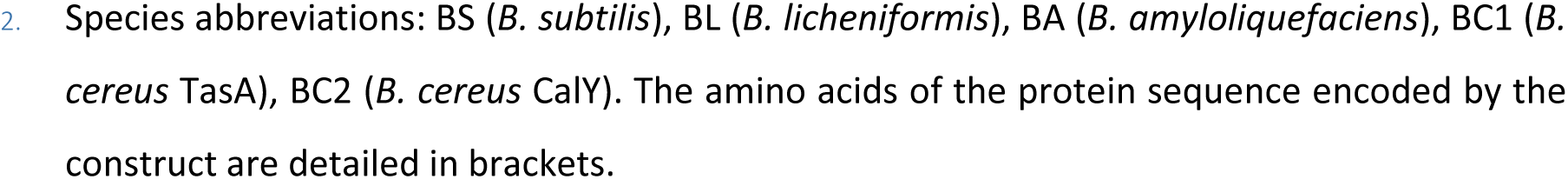
Plasmids used in this study

| Plasmid | Description | Source |
| --- | --- | --- |
| **pDR111** | *B. subtilis* integration vector for IPTG-induced | Britton *et al.*, 2002 |
| **pGEX-6P-1** | Vector for overexpression of GST-fused proteins | GE Healthcare |
| **pMiniMAD** | Temp sensitive allelic replacement vector | Patrick & Kearns., 2008 |
| **pNW1448** | pMinimad ΔtasA | This work |
| **pNW1619** | pDR110 $P_{IPTG}$-sipW-tasA (spc)<sup>1</sup> | This work |
| **pNW1437** | pDR110 $P_{IPTG}$-tasA (spc)<sup>1</sup> | This work |
| **pNW1631** | pDR110 $P_{IPTG}$-sipW<sub>TAA</sub>-tasA | This work |
| **pNW685** | pMAD ΔtapA | This work |
| **pNW1432** | pDR110 $P_{IPTG}$-tasA (spc)<sup>1</sup> | This work |
| **pNW1434** | pDR183 $P_{IPTG}$-tasA (erm)<sup>1</sup> | This work |
| **pNW543** | pGEX-6P-1- tasA<sub>(28-261)BS</sub> (amp)<sup>1,2</sup> | This work |
| **pNW1437** | pGEX-6P-1-TEV-tasA<sub>(28-261)BS</sub> (amp)<sup>1,2</sup> | This work |
| **pNW1080** | pGEX-6P-1-TEV-ser-tasA<sub>(28-261)BS</sub> (amp)<sup>1,2</sup> | This work |
| **pNW1082** | pGEX-6P-1-TEV-ala-tasA<sub>(28-261)BS</sub> (amp)<sup>1,2</sup> | This work |
| **pNW1083** | pGEX-6P-1-TEV-glu-tasA<sub>(28-261)BS</sub> (amp)<sup>1,2</sup> | This work |
| **pNW1084** | pGEX-6P-1-TEV-lys-tasA<sub>(28-261)BS</sub> (amp)<sup>1,2</sup> | This work |
| **pNW1085** | pGEX-6P-1-TEV-phe-tasA<sub>(28-261)BS</sub> (amp)<sup>1,2</sup> | This work |
| **pNW1606** | pGEX-6P-1-TEV-tasA<sub>(28-264)BL</sub> (amp)<sup>1,2</sup> | This work |
| **pNW1608** | pGEX-6P-1-TEV-tasA<sub>(28-261)BA</sub> (amp)<sup>1,2</sup> | This work |
| **pNW1096** | pGEX-6P-1-TEV-tasA<sub>(28-197)BC1</sub> (amp)<sup>1,2</sup> | This work |
| **pNW1616** | pGEX-6P-1-TEV-tasA<sub>(28-197)BC2</sub> (amp)<sup>1,2</sup> | This work |
| **pBL584** | epsA::pBL584Φ(Pspachy-epsA)-cat (cml)<sup>1</sup> | (Terra *et al.*, 2012) |
| **pNW1804** | pDR110- tapA-sipW-tasA | This work |

1. Drug resistance cassettes are indicated as follows: *spc*, spectinomycin, *amp*: ampicillin, *cml*: chloramphenicol, *erm*: erythromycin
2. Species abbreviations: BS (*B. subtilis*), BL (*B. licheniformis*), BA (*B. amyloliquefaciens*), BC1 (*B. cereus* TasA), BC2 (*B. cereus* CalY). The amino acids of the protein sequence encoded by the construct are detailed in brackets.

**Table S3.**
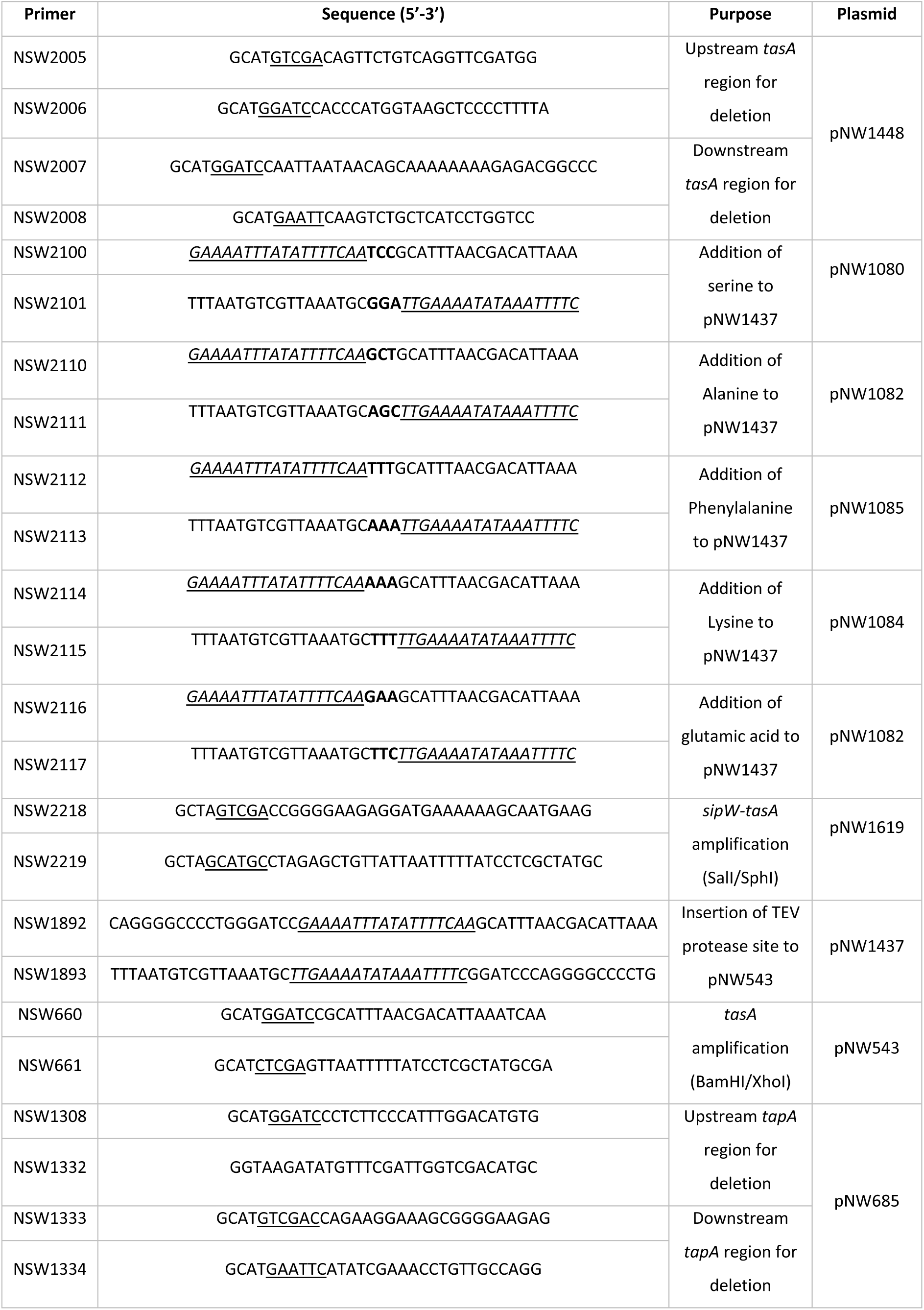
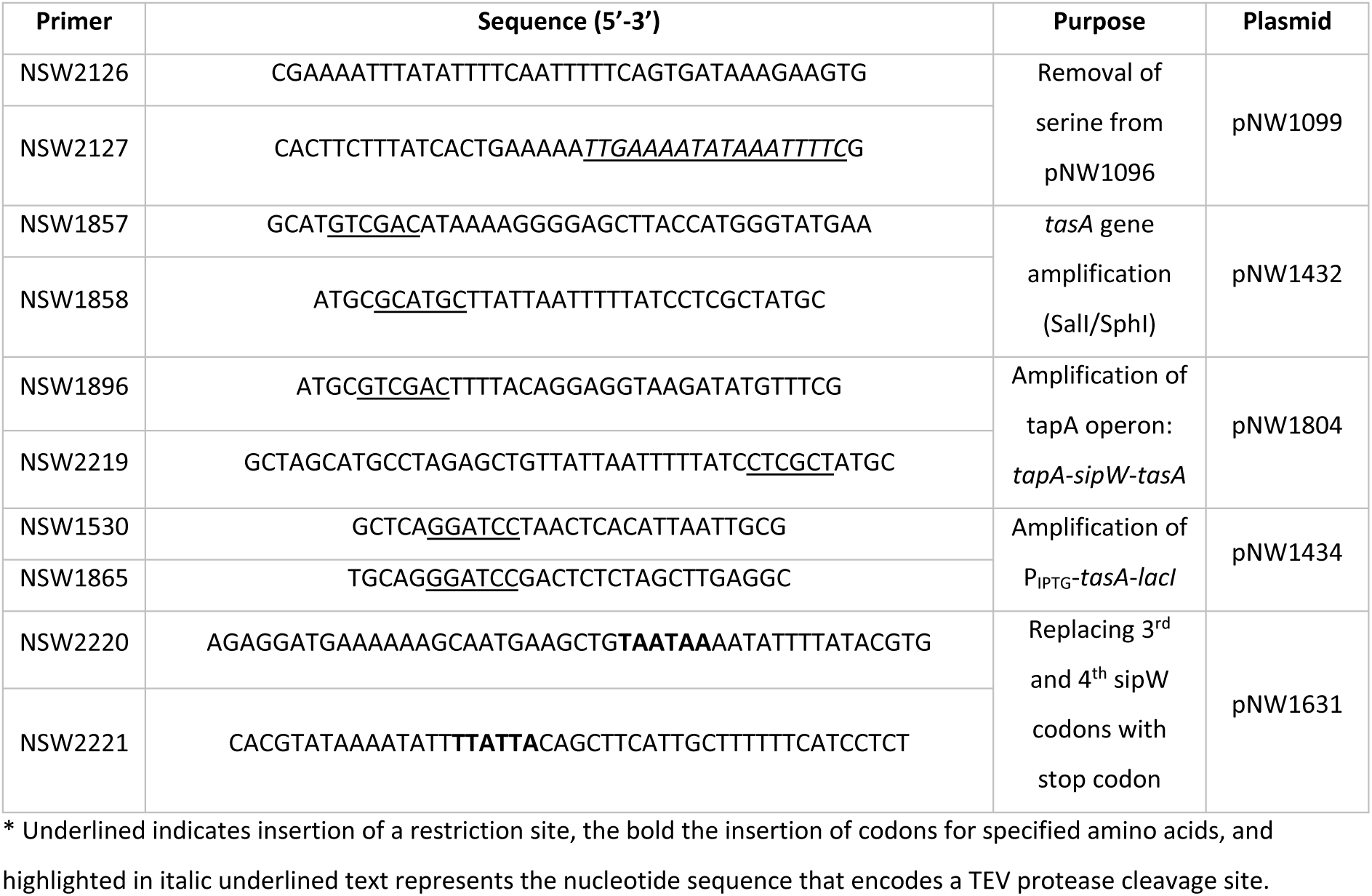
Primers used in this study.

**Table S4.**
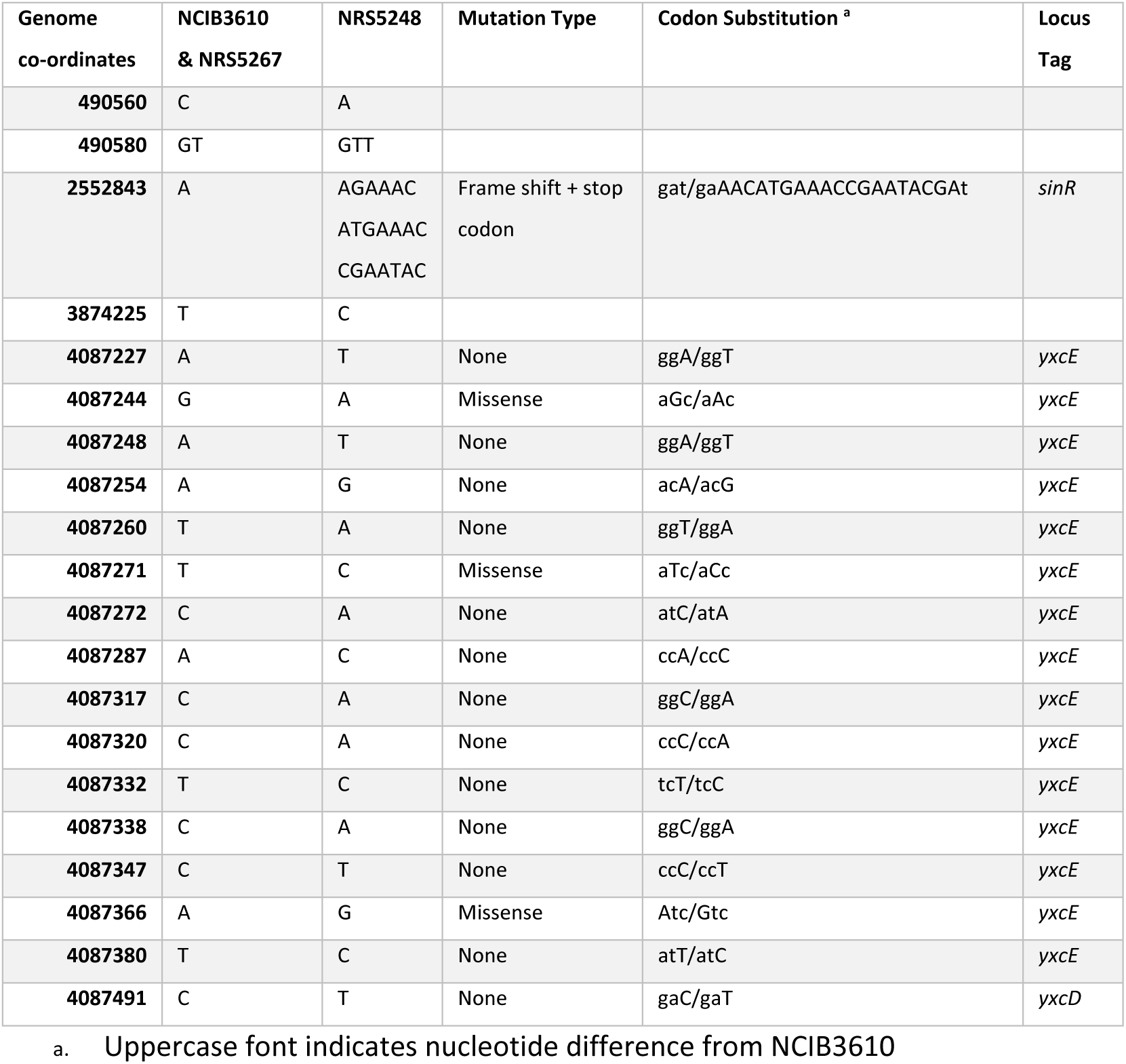
Single nucleotide polymorphisms identified by genomic sequencing

**Figure S1:**
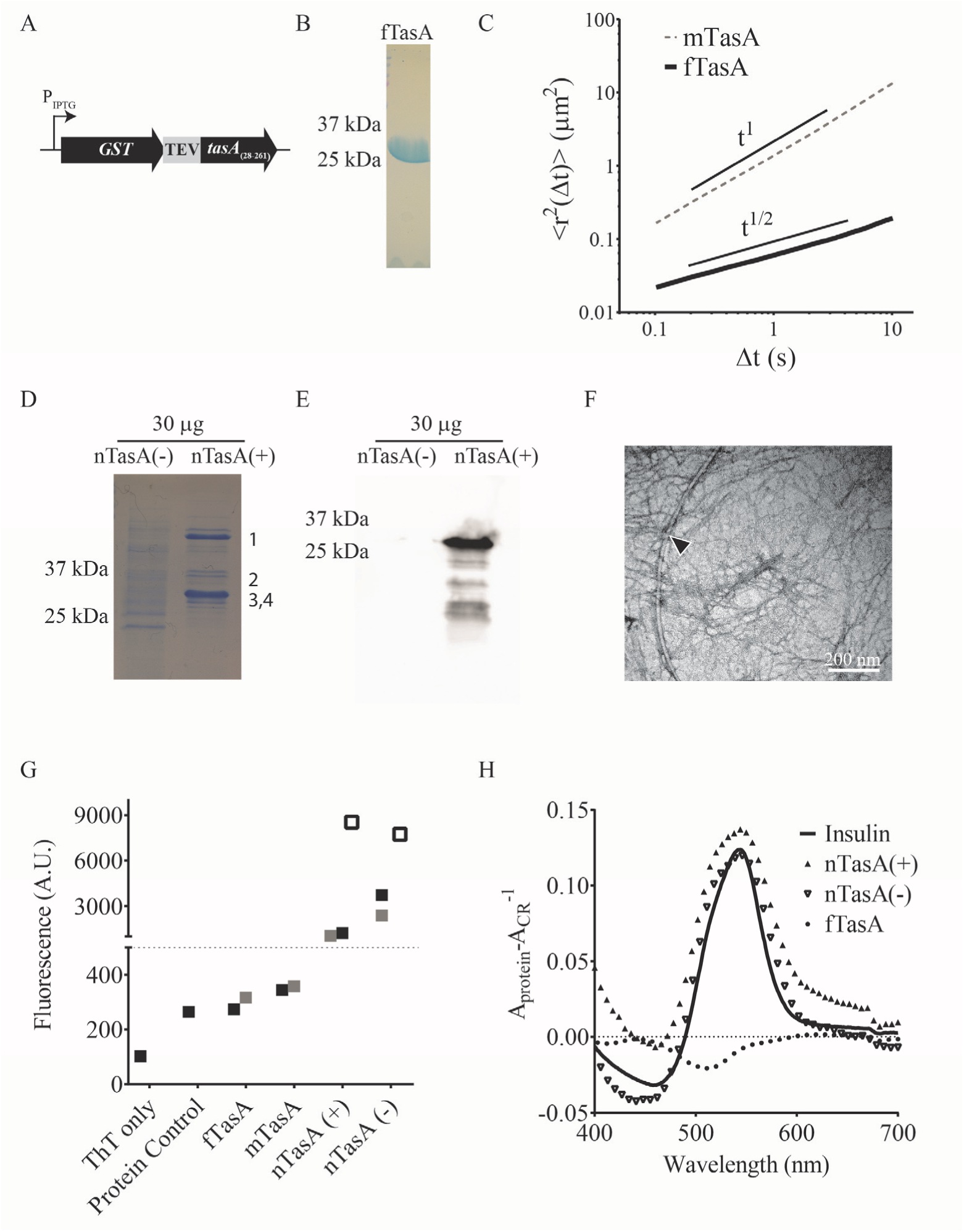
Purification and characterisation of recombinant fibre-forming TasA and natively extracted nTasA. (**A**) Schematic of expression construct used for purification of TasA(28-261) (fTasA) using the plasmid backbone pGex-6-P-1 with TEV protease cleavage site. (**B**) SDS-PAGE analysis of 10 μg purified fTasA with molecular mass of 25731 kDa as calculated by LC-MS analysis (**C**) Mean square displacement (MSD) versus lag time for 2 mg/ml fTasA (solid line) and mTasA (dashed line). The slope of the MSD for mTasA scales as t^1^ indicative of a viscous fluid medium. In contrast, the MSD slope for fTasA scales as t^1/2^ which is indicative of a viscoelastic medium. (**D**) SDS-PAGE analysis of 30 μg extracted nTasA(−) and nTasA(+) with bands 1, 2, 3 and 4 sent for identification by mass spectrometry. (**E**) Immunoblot analysis of 30 μg nTasA(−) and nTasA(+) extracts using anti-TasA antibody. (**F**) Transmission electron microscopy images of nTasA(+) stained with uranyl acetate with arrow highlighting flagella. (**G**) Maximum ThT fluorescence of ThT-only, protein control, nTasA(+) and nTasA(−) extracts showing 3 independent single reads. Recombinant fTasA and mTasA are the median read from 2 independent time course experiments. (**H**) Absorbance of Congo Red dye in presence of fTasA, nTasA(+) and nTasA(−) subtracted from background alongside insulin amyloid fibril positive control.

**Figure S2:**
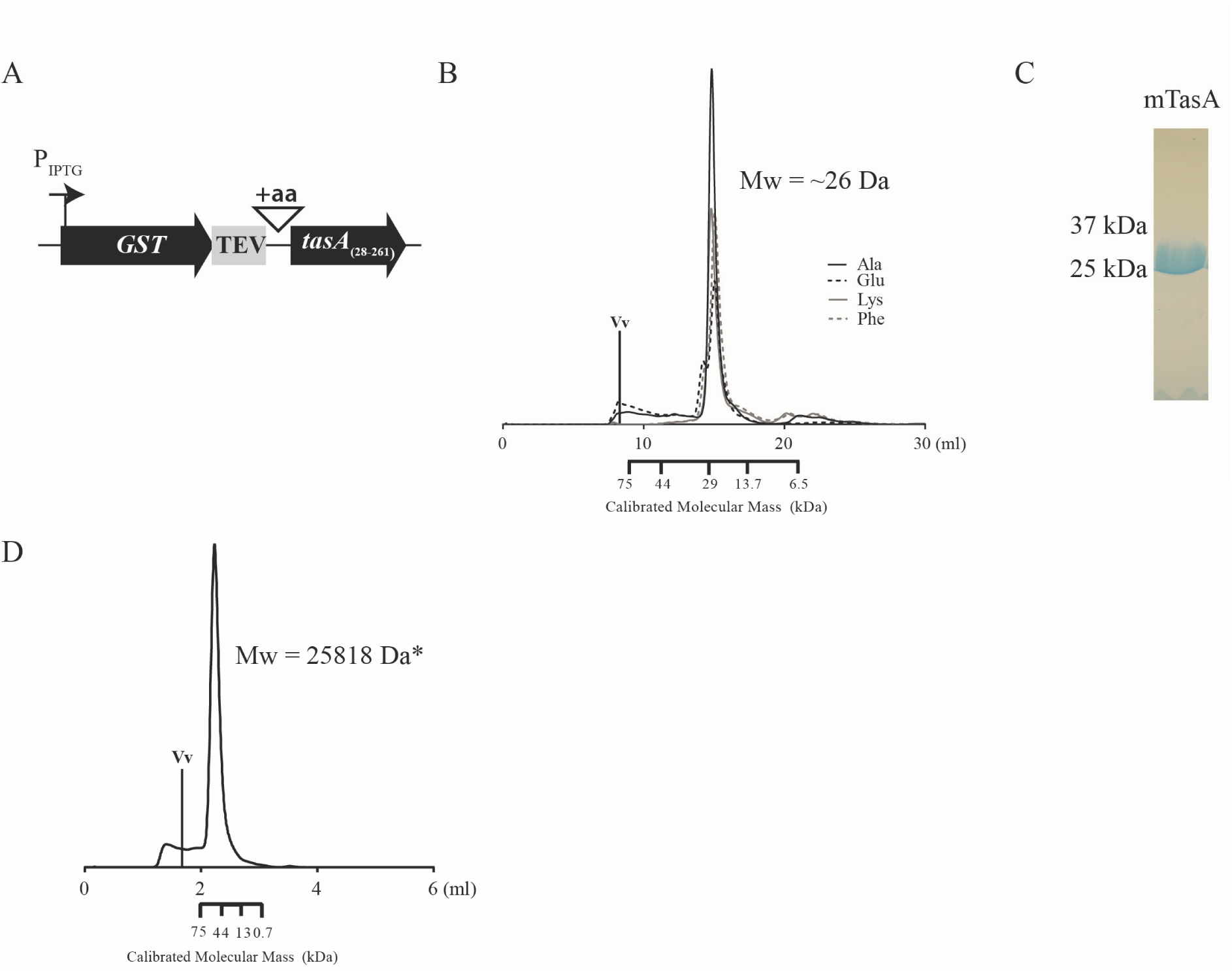
Purification and characterisation of recombinant monomeric TasA. (**A**) Sequence schematic of amino acid insertion, indicated by ‘aa’. (**B**) Size exclusion chromatography (SEC) analysis (Superdex 200 10/300 GL) of N-terminal tagged TasA where the amino acids at N-terminus indicated (Ala, Glu, Phe, Lys). (**C**) SDS-PAGE analysis of 10 μg purified serine tagged TasA (mTasA) with molecular mass of 25818 kDa as calculated by LC-MS. (**D**) SEC of mTasA (Superdex 200 5/150 GL).

**Figure S3:**
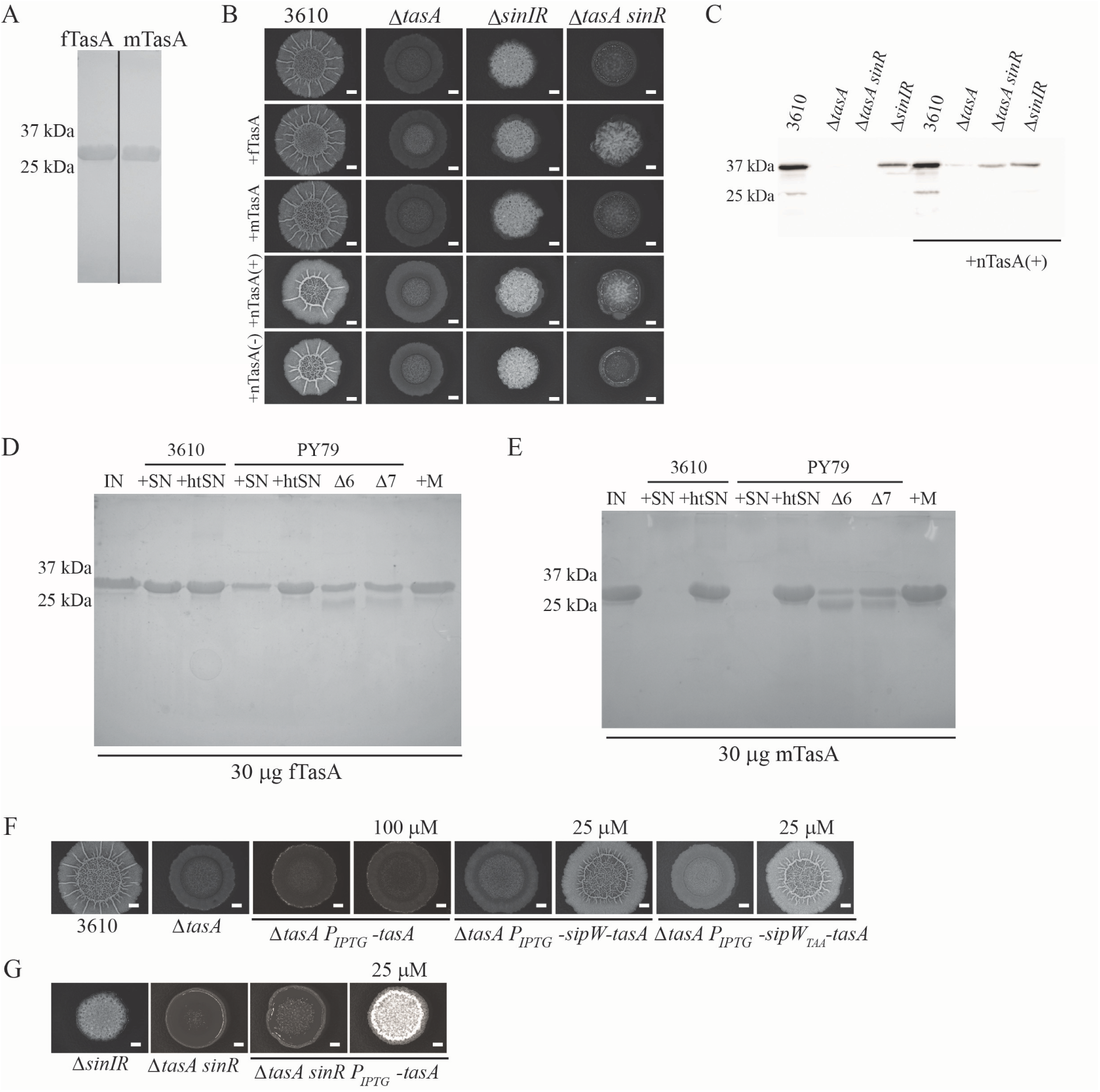
Recombinant fTasA is biologically active. (**A**) Representative SDS-PAGE analysis of fTasA and mTasA, showing 10 μg protein used in (B). (**B**) Biofilm phenotypes of NCIB3610, Δ*tasA* (NRS5267) *sinIR* (NRS2012) and Δ*tasA sinR* (NRS5248) with the *ex vivo* addition of 10 μg purified protein or 30 μg native extract as indicated. Images shown in Fig. 3A,C are repeated here. (**C**) Immunoblot blot analysis of biofilm lysate collected from controls and *ex vivo* addition of nTasA(+) challenged with α-TasA antibody. (**D-E**) Integrity of 30 μg fTasA and mTasA incubated for 24 hrs at 37°C analysed by SDS-PAGE. The protein (IN) was incubated with filtered spent supernatants collected from NCIB3610 and PY79 (+SN) and the same supernatants after heat inactivation at 100°C (+htSN); supernatants from exoprotease deficient strains derived from PY79 (Δ6 and Δ7) were also used. (+M) indicates media only control. (**F-G**) Control biofilms for the genetic complementation of Δ*tasA* and Δ*tasA sinR* as shown in Fig 2D-E in absence and presence of IPTG at concentrations indicated. Whole genome sequencing of Δ*tasA sinR* (NRS5248) strain identified a duplication of region Phe65-Iso71 leading to frame shift and stop codon as highlighted in Table S4.

**Figure S4:**
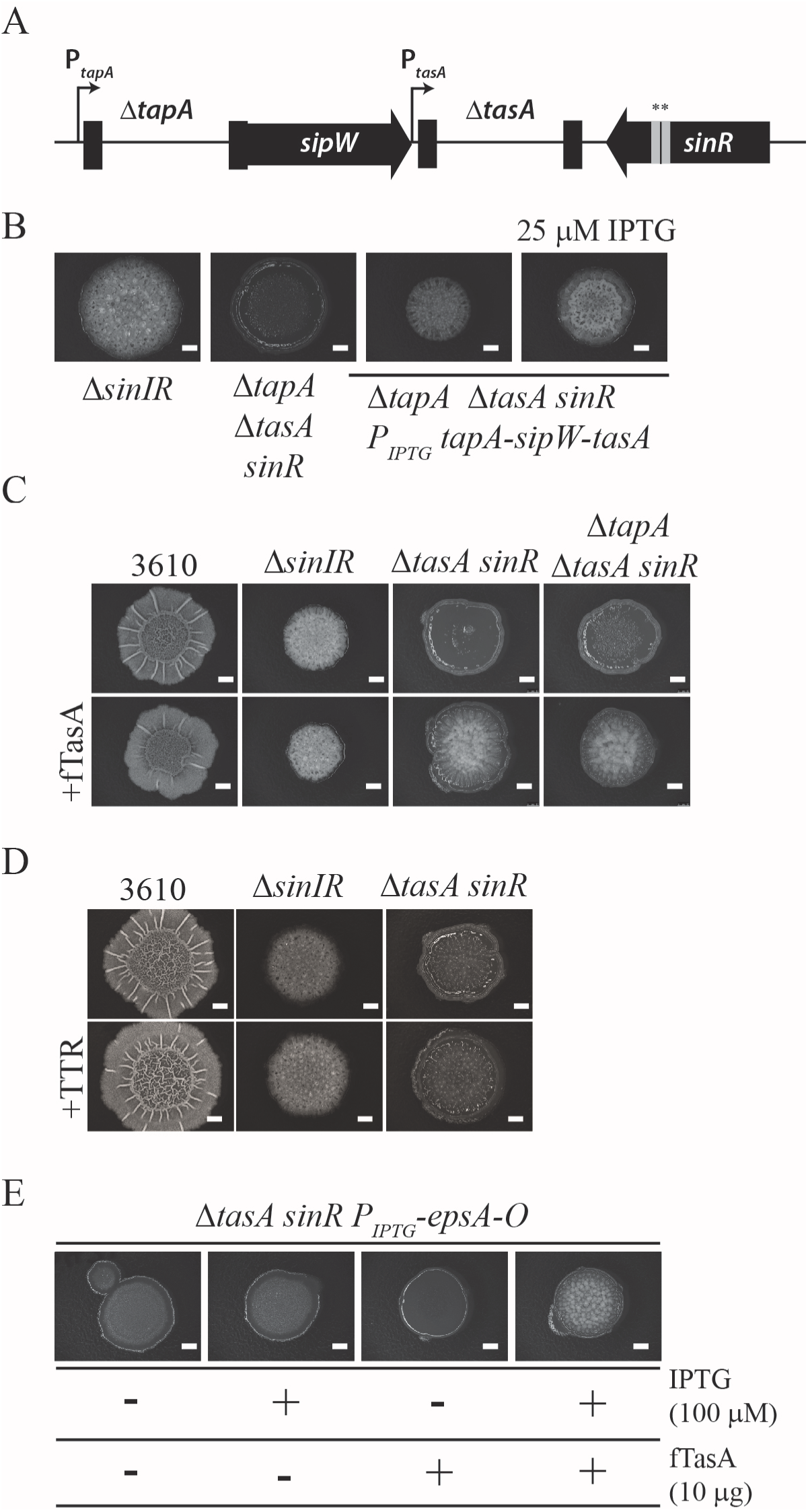
The biological activity of fTasA is independent of *tapA* but is dependent on specific matrix interaction. (**A**) Sequence schematic of Δ*tapA* Δ*tasA sinR* (NRS5749). (**B**) Biofilm phenotype of complementation in presence and absence of 25 μM IPTG (NRS5763). (**C**) Biofilm phenotypes of NCIB3610, Δ*tasA sinR* (NRS5248) and Δ*tapA* Δ*tasA sinR* (NRS5749) with the addition of 10 μg fTasA *ex vivo*. (**D**) Biofilm phenotype of Δ*tasA sinR* in presence and absence of the fibrous amyloid protein transthyretin (+TTR). (**E**) Biofilm phenotype of Δ*tasA sinR* P_iptg_-*eps* (NRS5421) in presence or absence of 100 μM IPTG and presence or absence of 10 μg fTasA where indicated by “−” and “+” respectively.

**Figure S5:**
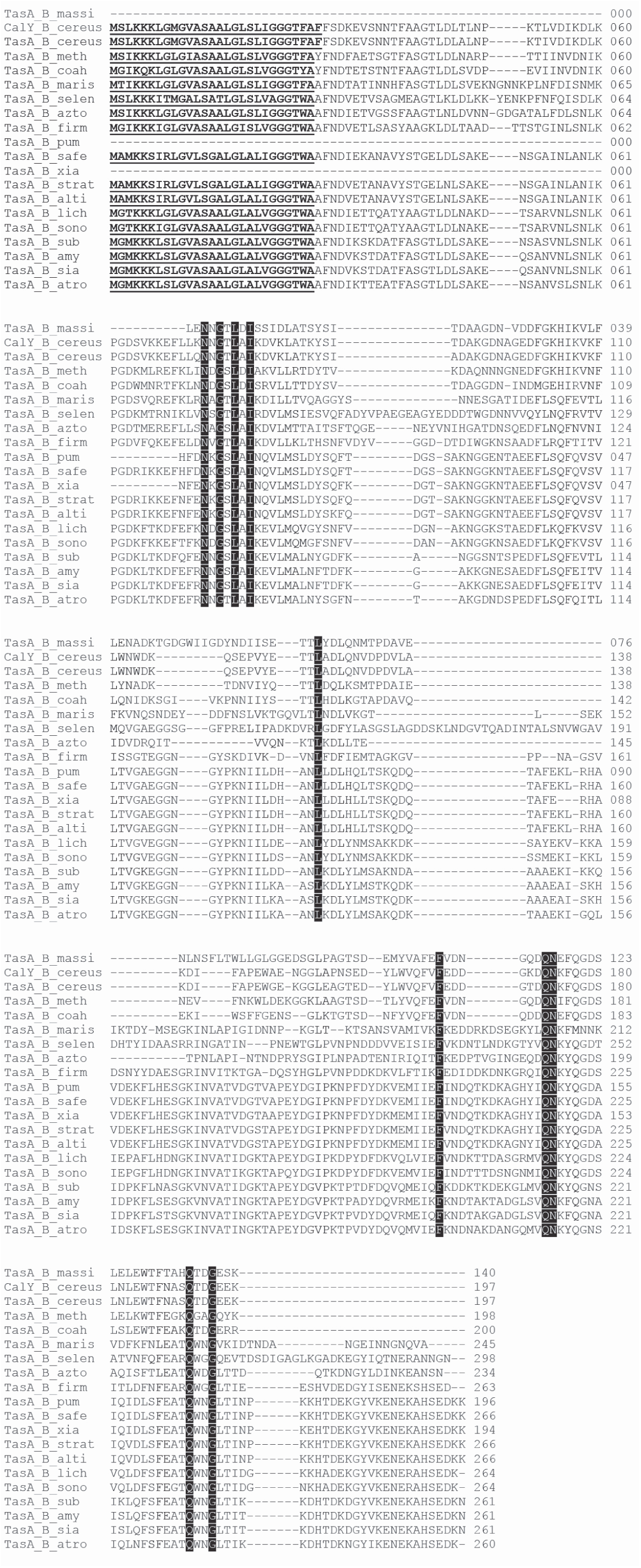
Alignment. Alignment of 20 TasA orthologues pulled from BlastP (Altschul *et al*., 1990; Altschul *et al*., 1997) of *B. subtilis* TasA sequence (*Bacillus azotoformans* (B_azto) TapA (WP_003329523), TasA (WP_035196521.1). *Bacillus firmus* (B_firm) TapA (WP_082139007), TasA (WP_035326610.1). *Bacillus selenatarsenatis* (B_selen) TapA (WP_084135527), TasA (WP_041967097.1). *Bacillus licheniformis* (B_lich) TapA (WP_075747486), TasA (WP_043927876.1). *Bacillus sonorensis* (B_sono) TapA (WP_006637531), TasA (WP_006637529.1). *Bacillus marisflavi* (B_maris) TapA (WP_082139781), TasA (WP_048012492.1). *Bacillus amyloliquefaciens* (B_amy) TapA (WP_063094776), TasA (WP_044802563.1). *Bacillus siamensis* (B_sia) TapA (WP_029575370), TasA (WP_045926714.1). *Bacillus atrophaeus* (B_atro) TapA (WP_061670003), TasA (WP_010789194.1). *Bacillus stratosphericus* (B_strat) TapA (WP_039964022), TasA WP (007501219.1). *Bacillus altitudinis* (B_alti) TapA (WP_073413951), TasA (WP_039166017.1). *Bacillus xiamenensis* (B_xia) TapA (WP_034739525), TasA (WP_008360245.1). *Bacillus pumilis* (B_pum) TapA (WP_041106857), TasA (WP_041106853.1). *Bacillus safensis* (B_safe) TapA (WP_075623495), TasA (WP_034280222.1). *Bacillus methanolicus* (B_meth) TapA (None), TasA (WP_004434174.1). *Bacillus massiliosenegalensis* (B_massi) TapA (None), TasA (WP_019154446.1). *Bacillus coahuilensis* (B_coah) TapA (None), TasA (WP_010174983.1). *Bacillus cereus* (B_cer) TapA (None), TasA (WP_002201283.1)) by Clustal Omega (Sievers *et al*., 2011). Signal sequence as predicted by SignalP v4.2 (Petersen *et al*., 2011) underlined, 100% sequence identity highlighted in black.

**Figure S6:**
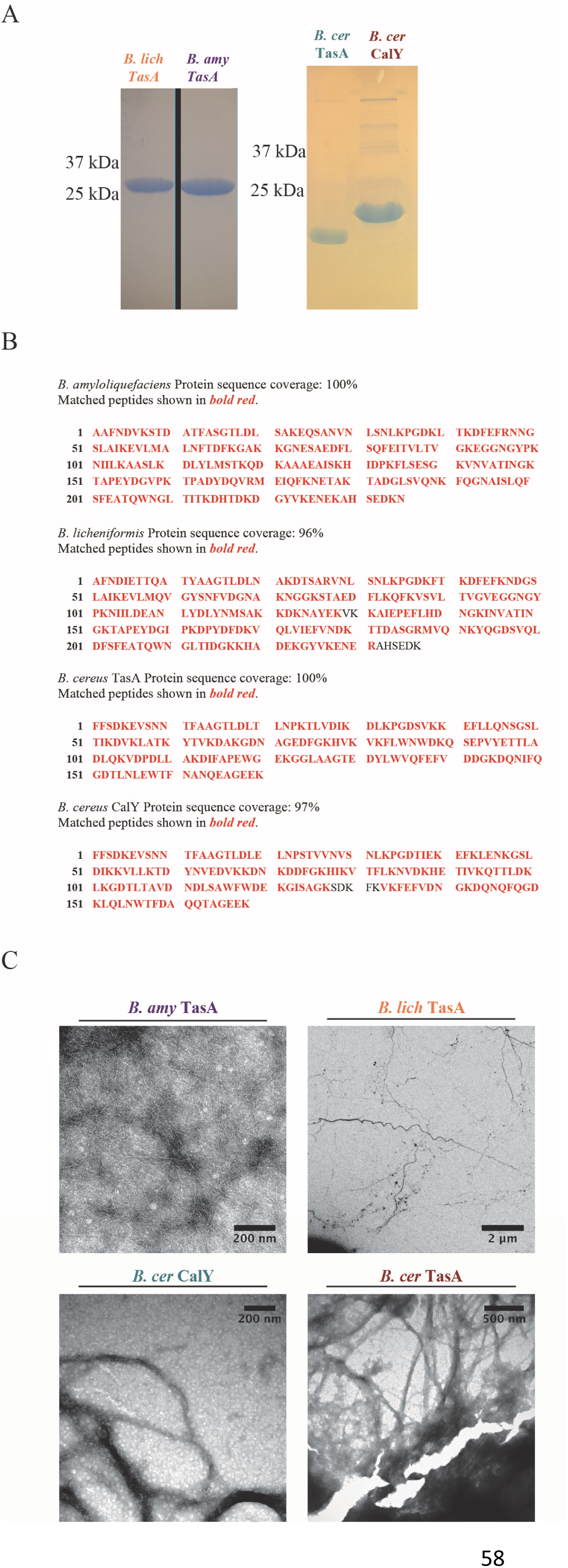
Characterisation of recombinant orthologous TasA. (**A**) SDS-PAGE of recombinant 10 μg *B. licheniformis, B. amyloliquefaciens* and *B. cereus* TasA and CalY generated in *E. coli*. (**B**) Coverage map of orthologue recombinant proteins sequences as determined by tandem mass spectrometry. Amino acid designated 1 signifies the first amino acid of the mature protein: *B. amyloliquefaciens* amino acid 27, *B. licheniformis* amino acid 28 and *B. cereus* for both CalY and TasA amino acid 30. (**C**) Transmission electron microscopy images of recombinant orthologue TasA stained with uranyl acetate show presence of fibres which are several micrometres long and approximately 22nm wide, with *B. cereus* TasA fibres being significantly wider at approximately 60nm.

**Figure S7:**
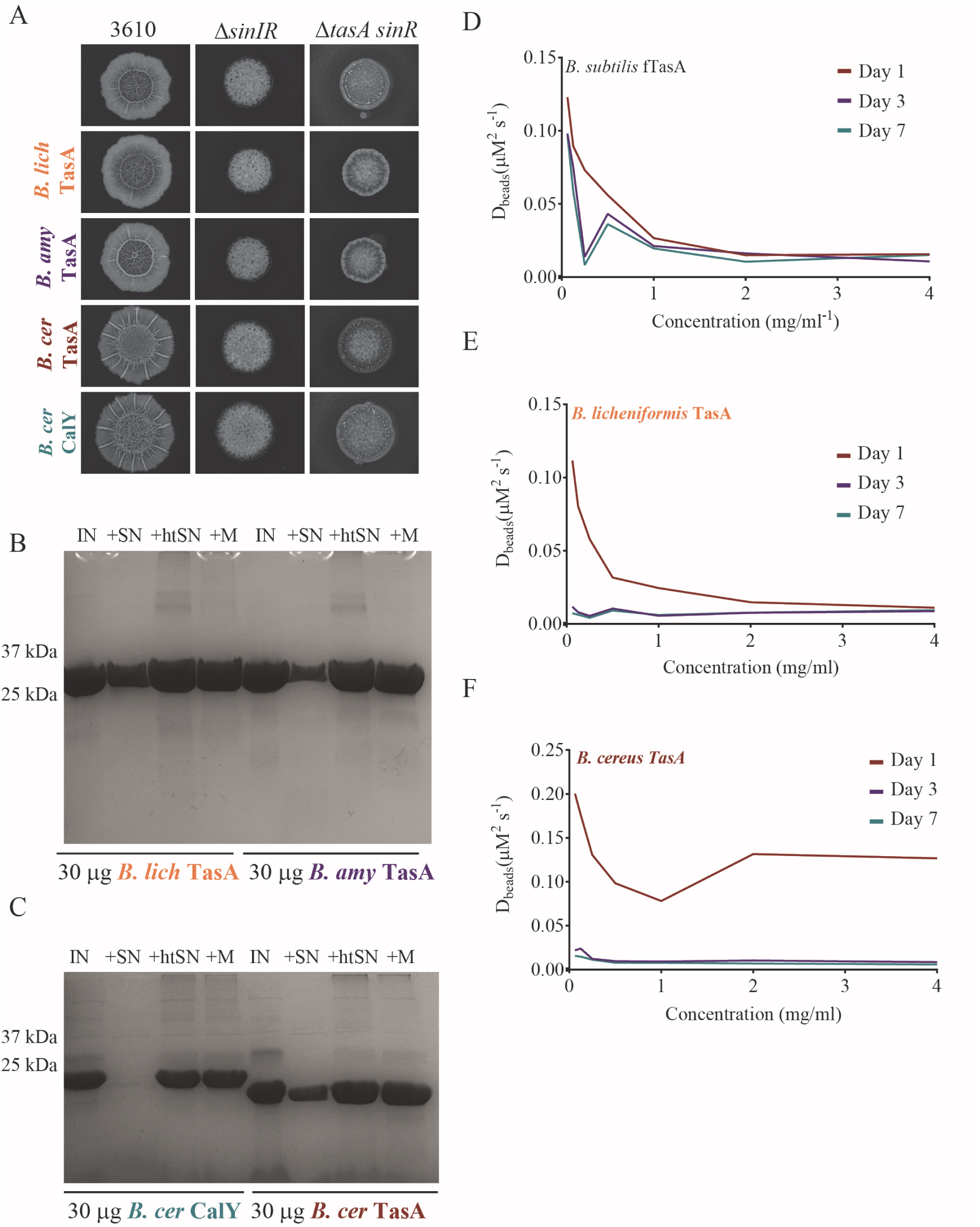
Complementation of Δ*tasA sinR* by recombinant orthologous fTasA. (**A**) Biofilm phenotypes of wild type (NCIB3610), Δ*tasA* (NRS5267), Δ*sinIR* (NRS2012) and Δ*tasA sinR* (NRS5248) strains with *ex vivo* addition of 10 μg recombinant orthologous TasA as indicated. (**B-C**) Integrity of 30 μg *B. licheniformis* TasA, *B. amyloliquefaciens* TasA and *B. cereus* CalY and TasA incubated for 24 hrs at 37°C then analysed by SDS-PAGE. The protein (IN) was incubated with filtered supernatant collected from NCIB3610 (+SN) and the supernatant after heat inactivation at 100°C (+htSN) alongside a media only control (+M). (**D-F**) Serial dilutions (from 4 mg/ml down to 0.0625 mg/ml) were prepared for *B. subtilis, B. licheniformis*, and *B. cereus* fTasA. Diffusion coefficients of 1μm beads were extracted from mean squared displacement curves. The diffusion coefficients were measured over successive days. The *B. licheniformis* and *B. cereus* fTasA samples gelled at lower concentrations compared to *B. subtilis* fTasA. Moreover, after 3 days, the *B. licheniformis* and *B. cereus* fTasA samples became gels for all concentrations studied indicating a slow, dynamical gelling process.

## References

Altschul, S.F., Gish, W., Miller, W., Myers, E.W., and Lipman, D.J. (1990) Basic local alignment search tool. J Mol Biol 215: 403–410

Altschul, S.F., Madden, T.L., Schäffer, A.A., Zhang, J., Zhang, Z., Miller, W., and Lipman, D.J. (1997) Gapped BLAST and PSI-BLAST: a new generation of protein database search programs. Nucleic Acids Res 25: 3389–402 http://www.ncbi.nlm.nih.gov/pubmed/9254694. Accessed August 31, 2017.

Arnaouteli, S., Ferreira, A.S., Schor, M., Morris, R.J., Bromley, K.M., Jo, J., et al. (2017) Bifunctionality of a biofilm matrix protein controlled by redox state. Proc Natl Acad Sci 114: E6184–E6191 http://www.ncbi.nlm.nih.gov/pubmed/28698374. Accessed August 31, 2017.

Arnaud, M., Chastanet, A., and Débarbouillé, M. (2004) New vector for efficient allelic replacement in naturally nontransformable, low-GC-content, gram-positive bacteria. Appl Environ Microbiol 70: 6887–91 http://www.pubmedcentral.nih.gov/articlerender.fcgi?artid=525206&tool=pmcentrez&rendertype=abstract. Accessed June 22, 2015.

Assefa, S., Keane, T.M., Otto, T.D., Newbold, C., and Berriman, M. (2009) ABACAS: algorithm-based automatic contiguation of assembled sequences. Bioinformatics 25: 1968–9 http://www.ncbi.nlm.nih.gov/pubmed/19497936. Accessed July 20, 2016.

Bais, H.P., Fall, R., and Vivanco, J.M. (2004) Biocontrol of Bacillus subtilis against infection of Arabidopsis roots by Pseudomonas syringae is facilitated by biofilm formation and surfactin production. Plant Physiol 134: 307–19 http://www.pubmedcentral.nih.gov/articlerender.fcgi?artid=316310&tool=pmcentrez&rendertype=abstract. Accessed December 15, 2014.

Baldwin, A.J., Knowles, T.P.J., Tartaglia, G.G., Fitzpatrick, A.W., Devlin, G.L., Shammas, S.L., et al. (2011) Metastability of Native Proteins and the Phenomenon of Amyloid Formation. J Am Chem Soc 133: 14160–14163 http://www.ncbi.nlm.nih.gov/pubmed/21650202. Accessed August 30, 2017.

Bankevich, A., Nurk, S., Antipov, D., Gurevich, A.A., Dvorkin, M., Kulikov, A.S., et al. (2012) SPAdes: a new genome assembly algorithm and its applications to single-cell sequencing. J Comput Biol 19: 455–77 http://www.ncbi.nlm.nih.gov/pubmed/22506599. Accessed July 20, 2016.

Beauregard, P.B., Chai, Y., Vlamakis, H., Losick, R., and Kolter, R. (2013) Bacillus subtilis biofilm induction by plant polysaccharides. Proc Natl Acad Sci U S A 110: E1621–30 http://www.ncbi.nlm.nih.gov/pubmed/23569226. Accessed September 5, 2017.

Benditt, E.P. (1986) Amyloid Protein AA and its Precursor, The Acute Phase Protein(s) ApoSAA: A Perspective. In Amyloidosis. Marrink, J., and M.H. van Rijswijk (eds). Springer Netherlands, Dordrecht. pp. 101–106 http://link.springer.com/10.1007/978-94-009-4309-4_11. Accessed September 5, 2017.

Bolger, A.M., Lohse, M., and Usadel, B. (2014) Trimmomatic: a flexible trimmer for Illumina sequence data. Bioinformatics 30: 2114–20 http://www.ncbi.nlm.nih.gov/pubmed/24695404. Accessed July 20, 2016.

Branda, S.S., Chu, F., Kearns, D.B., Losick, R., and Kolter, R. (2006) A major protein component of the Bacillus subtilis biofilm matrix. Mol Microbiol 59: 1229–38 http://www.ncbi.nlm.nih.gov/pubmed/16430696. Accessed November 3, 2014.

Branda, S.S., González-Pastor, J.E., Ben-Yehuda, S., Losick, R., and Kolter, R. (2001) Fruiting body formation by Bacillus subtilis. Proc Natl Acad Sci U S A 98: 11621–6 http://www.pubmedcentral.nih.gov/articlerender.fcgi?artid=58779&tool=pmcentrez&rendertype=abstract. Accessed May 15, 2016.

Branda, S.S., Gonzalez-Pastor, J.E., Dervyn, E., Ehrlich, S.D., Losick, R., and Kolter, R. (2004) Genes Involved in Formation of Structured Multicellular Communities by Bacillus subtilis. J Bacteriol 186: 3970–3979 http://www.ncbi.nlm.nih.gov/pubmed/15175311. Accessed April 24, 2017.

Bromley, K.M., Morris, R.J., Hobley, L., Brandani, G., Gillespie, R.M.C., McCluskey, M., et al. (2015) Interfacial self-assembly of a bacterial hydrophobin. Proc Natl Acad Sci U S A 112: 5419–24 http://www.ncbi.nlm.nih.gov/pubmed/25870300. Accessed August 11, 2017.

Cairns, L.S., Hobley, L., and Stanley-Wall, N.R. (2014) Biofilm formation by Bacillus subtilis: new insights into regulatory strategies and assembly mechanisms. Mol Microbiol 93: 587–98 http://www.pubmedcentral.nih.gov/articlerender.fcgi?artid=4238804&tool=pmcentrez&rendertype=abstract. Accessed July 10, 2014.

Caro-Astorga, J., PÃ©rez-GarcÃ-a, A., Vicente, A. de, and Romero, D. (2015) A genomic region involved in the formation of adhesin fibers in Bacillus cereus biofilms. Front Microbiol 5: 1–11 http://journal.frontiersin.org/article/10.3389/fmicb.2014.00745/abstract.

Castresana, J. (2000) Selection of conserved blocks from multiple alignments for their use in phylogenetic analysis. Mol Biol Evol 17: 540–52 http://www.ncbi.nlm.nih.gov/pubmed/10742046. Accessed May 19, 2017.

Chai, L., Romero, D., Kayatekin, C., Akabayov, B., Vlamakis, H., Losick, R., and Kolter, R. (2013) Isolation, characterization, and aggregation of a structured bacterial matrix precursor. J Biol Chem 288: 17559–68 http://www.ncbi.nlm.nih.gov/pubmed/23632024. Accessed June 1, 2017.

Chai, Y., Beauregard, P.B., Vlamakis, H., Losick, R., and Kolter, R. (2012) Galactose Metabolism Plays a Crucial Role in Biofilm Formation by Bacillus subtilis. MBio 3: e00184-12-e00184-12 http://www.ncbi.nlm.nih.gov/pubmed/22893383. Accessed September 5, 2017.

Chapman, M.R., Robinson, L.S., Pinkner, J.S., Roth, R., Heuser, J., Hammar, M., et al. (2002) Role of Escherichia coli curli operons in directing amyloid fiber formation. Science 295: 851–5 http://www.ncbi.nlm.nih.gov/pubmed/11823641. Accessed January 17, 2018.

Chen, Y., Cao, S., Chai, Y., Clardy, J., Kolter, R., Guo, J., and Losick, R. (2012) A Bacillus subtilis sensor kinase involved in triggering biofilm formation on the roots of tomato plants. Mol Microbiol 85: 418–30 http://www.ncbi.nlm.nih.gov/pubmed/22716461. Accessed April 25, 2017.

Chen, Y., Yan, F., Chai, Y., Liu, H., Kolter, R., Losick, R., and Guo, J. (2013) Biocontrol of tomato wilt disease by Bacillus subtilis isolates from natural environments depends on conserved genes mediating biofilm formation. Environ Microbiol 15: 848–64 http://www.ncbi.nlm.nih.gov/pubmed/22934631. Accessed April 19, 2017.

Chevenet, F., Brun, C., Bañuls, A.-L., Jacq, B., and Christen, R. (2006) TreeDyn: towards dynamic graphics and annotations for analyses of trees. BMC Bioinformatics 7: 439 http://www.ncbi.nlm.nih.gov/pubmed/17032440. Accessed May 19, 2017.

Chu, F., Kearns, D.B., Branda, S.S., Kolter, R., and Losick, R. (2006) Targets of the master regulator of biofilm formation in Bacillus subtilis. Mol Microbiol 59: 1216–1228 http://www.ncbi.nlm.nih.gov/pubmed/16430695. Accessed October 10, 2015.

Cianfanelli, F.R., Alcoforado Diniz, J., Guo, M., Cesare, V. De, Trost, M., Coulthurst, S.J., et al. (2016) VgrG and PAAR Proteins Define Distinct Versions of a Functional Type VI Secretion System. PLOS Pathog 12: e1005735 http://dx.plos.org/10.1371/journal.ppat.1005735. Accessed August 9, 2016.

Costerton, J.W., Cheng, K.J., Geesey, G.G., Ladd, T.I., Nickel, J.C., Dasgupta, M., and Marrie, T.J. (1987) Bacterial Biofilms in Nature and Disease. Annu Rev Microbiol 41: 435–464 http://www.annualreviews.org/doi/10.1146/annurev.mi.41.100187.002251. Accessed August 22, 2016.

Dereeper, A., Guignon, V., Blanc, G., Audic, S., Buffet, S., Chevenet, F., et al. (2008) Phylogeny.fr: robust phylogenetic analysis for the non-specialist. Nucleic Acids Res 36: W465–W469 http://www.ncbi.nlm.nih.gov/pubmed/18424797. Accessed May 19, 2017.

Dobson, C.M. (1999) Protein misfolding, evolution and disease. Trends Biochem Sci 24: 329–332 http://linkinghub.elsevier.com/retrieve/pii/S0968000499014450. Accessed August 11, 2017.

Dragoš, A., and Kovács, Á.T. (2017) The Peculiar Functions of the Bacterial Extracellular Matrix. Trends Microbiol 25: 257–266 http://www.ncbi.nlm.nih.gov/pubmed/28089324. Accessed August 31, 2017.

Edelstein, A., Amodaj, N., Hoover, K., Vale, R., and Stuurman, N. (2010) Computer control of microscopes using μManager. Curr Protoc Mol Biol **Chapter 14**: Unit14.20 http://www.ncbi.nlm.nih.gov/pubmed/20890901. Accessed August 30, 2017.

Eisenberg, D., and Jucker, M. (2012) The amyloid state of proteins in human diseases. Cell 148: 1188–203 http://www.ncbi.nlm.nih.gov/pubmed/22424229. Accessed August 11, 2017.

Elsholz, A.K.W., Wacker, S.A., and Losick, R. (2014) Self-regulation of exopolysaccharide production in Bacillus subtilis by a tyrosine kinase. Genes Dev 28: 1710–1720 http://www.ncbi.nlm.nih.gov/pubmed/25085422. Accessed September 1, 2017.

Epstein, E.A., and Chapman, M.R. (2008) Polymerizing the fibre between bacteria and host cells: the biogenesis of functional amyloid fibres. Cell Microbiol 10: 1413–20 http://www.ncbi.nlm.nih.gov/pubmed/18373633. Accessed January 14, 2018.

Evans, M.L., and Chapman, M.R. (2014) Curli biogenesis: order out of disorder. Biochim Biophys Acta 1843: 1551–8 http://www.ncbi.nlm.nih.gov/pubmed/24080089. Accessed January 13, 2018.

Ferrão-Gonzales, A.D., Souto, S.O., Silva, J.L., and Foguel, D. (2000) The preaggregated state of an amyloidogenic protein: hydrostatic pressure converts native transthyretin into the amyloidogenic state. Proc Natl Acad Sci U S A 97: 6445–50 http://www.ncbi.nlm.nih.gov/pubmed/10841549. Accessed September 5, 2017.

Ferrari, G. V De, Mallender, W.D., Inestrosa, N.C., and Rosenberry, T.L. (2001) Thioflavin T is a fluorescent probe of the acetylcholinesterase peripheral site that reveals conformational interactions between the peripheral and acylation sites. J Biol Chem 276: 23282–7 http://www.ncbi.nlm.nih.gov/pubmed/11313335. Accessed August 30, 2017.

Fitzpatrick, A.W.P., Debelouchina, G.T., Bayro, M.J., Clare, D.K., Caporini, M.A., Bajaj, V.S., et al. (2013) Atomic structure and hierarchical assembly of a cross-ß amyloid fibril. Proc Natl Acad Sci U S A 110: 5468–73 http://www.ncbi.nlm.nih.gov/pubmed/23513222. Accessed September 5, 2017.

Fitzpatrick, A.W.P., Falcon, B., He, S., Murzin, A.G., Murshudov, G., Garringer, H.J., et al. (2017) Cryo-EM structures of tau filaments from Alzheimer’s disease. Nat Publ Gr 547 https://www.nature.com/nature/journal/v547/n7662/pdf/nature23002.pdf. Accessed August 11, 2017.

Flemming, H.-C., and Wingender, J. (2010) The biofilm matrix. Nat Rev Microbiol 8: 623 http://www.nature.com/doifinder/10.1038/nrmicro2415. Accessed August 31, 2017.

Fowler, D.M., Koulov, A. V., Balch, W.E., and Kelly, J.W. (2007) Functional amyloid - from bacteria to humans. Trends Biochem Sci 32: 217–224 http://www.ncbi.nlm.nih.gov/pubmed/17412596. Accessed August 11, 2017.

Freire, S., Araujo, M.H. de, Al-Soufi, W., and Novo, M. (2014) Photophysical study of Thioflavin T as fluorescence marker of amyloid fibrils. Dye Pigment 110: 97–105.

Gestel, J. van, Vlamakis, H., and Kolter, R. (2015) From Cell Differentiation to Cell Collectives: Bacillus subtilis Uses Division of Labor to Migrate. PLOS Biol 13: e1002141 http://journals.plos.org/plosbiology/article?id=10.1371/journal.pbio.1002141. Accessed April 21, 2015.

Guindon, S., and Gascuel, O. (2003) A simple, fast, and accurate algorithm to estimate large phylogenies by maximum likelihood. Syst Biol 52: 696–704 http://www.ncbi.nlm.nih.gov/pubmed/14530136. Accessed May 19, 2017.

Hardy, J., and Selkoe, D.J. (2002) The Amyloid Hypothesis of Alzheimer’s Disease: Progress and Problems on the Road to Therapeutics. Science (80-) 297: 353–356 http://www.ncbi.nlm.nih.gov/pubmed/12130773. Accessed September 5, 2017.

Hobley, L., Harkins, C., MacPhee, C.E., and Stanley-Wall, N.R. (2015) Giving structure to the biofilm matrix: an overview of individual strategies and emerging common themes. FEMS Microbiol Rev fuv015 http://femsre.oxfordjournals.org/content/early/2015/04/22/femsre.fuv015.abstract. Accessed April 24, 2015.

Hobley, L., Ostrowski, A., Rao, F. V, Bromley, K.M., Porter, M., Prescott, A.R., et al. (2013) BslA is a self-assembling bacterial hydrophobin that coats the Bacillus subtilis biofilm. Proc Natl Acad Sci U S A 110: 13600–5 http://www.pubmedcentral.nih.gov/articlerender.fcgi?artid=3746881&tool=pmcentrez&rendertype=abstract. Accessed October 31, 2014.

Hong, D.-P., Ahmad, A., and Fink, A.L. (2006) Fibrillation of Human Insulin A and B Chains. Biochemistry 45: 9342–9353 http://www.ncbi.nlm.nih.gov/pubmed/16866381. Accessed September 5, 2017.

Howie, A.J., and Brewer, D.B. (2009) Optical properties of amyloid stained by Congo red: History and mechanisms. Micron 40: 285–301 https://www.sciencedirect.com/science/article/pii/S0968432808002187?via%3Dihub. Accessed January 24, 2018.

Jones, S.E., Paynich, M.L., Kearns, D.B., and Knight, K.L. (2014) Protection from Intestinal Inflammation by Bacterial Exopolysaccharides. J Immunol 192: 4813–4820 http://www.ncbi.nlm.nih.gov/pubmed/24740503. Accessed September 5, 2017.

Kajava, A. V., and Steven, A.C. (2006) ß-Rolls, ß-Helices, and Other β-Solenoid Proteins. In Advances in protein chemistry. pp. 55–96 http://www.ncbi.nlm.nih.gov/pubmed/17190611. Accessed September 5, 2017.

Kalapothakis, J.M.D., Morris, R.J., Szavits-Nossan, J., Eden, K., Covill, S., Tabor, S., et al. (2015) A kinetic study of ovalbumin fibril formation: the importance of fragmentation and end-joining. Biophys J 108: 2300–11 http://www.ncbi.nlm.nih.gov/pubmed/25954887. Accessed September 5, 2017.

Kayed, R., Bernhagen, J., Greenfield, N., Sweimeh, K., Brunner, H., Voelter, W., and Kapurniotu, A. (1999) Conformational transitions of islet amyloid polypeptide (IAPP) in amyloid formation in Vitro. J Mol Biol 287: 781–796 http://www.ncbi.nlm.nih.gov/pubmed/10191146. Accessed September 5, 2017.

Kiley, T.B., and Stanley-Wall, N.R. (2010) Post-translational control of Bacillus subtilis biofilm formation mediated by tyrosine phosphorylation. Mol Microbiol 78: 947–963 http://www.ncbi.nlm.nih.gov/pubmed/20815827. Accessed April 19, 2017.

Koboldt, D.C., Chen, K., Wylie, T., Larson, D.E., McLellan, M.D., Mardis, E.R., et al. (2009) VarScan: variant detection in massively parallel sequencing of individual and pooled samples. Bioinformatics 25: 2283–5 http://www.ncbi.nlm.nih.gov/pubmed/19542151. Accessed July 20, 2016.

LeVine, H. (1999) [18] Quantification of β-sheet amyloid fibril structures with thioflavin T. Methods Enzymol 309: 274–284.

Li, H., Handsaker, B., Wysoker, A., Fennell, T., Ruan, J., Homer, N., et al. (2009) The Sequence Alignment/Map format and SAMtools. Bioinformatics 25: 2078–9 http://www.ncbi.nlm.nih.gov/pubmed/19505943. Accessed July 20, 2016.

MacPhee, C.E., and Dobson, C.M. (2000) Formation of Mixed Fibrils Demonstrates the Generic Nature and Potential Utility of Amyloid Nanostructures. http://pubs.acs.org/doi/full/10.1021/ja0029580. Accessed September 5, 2017.

Makin, O.S., and Serpell, L.C. (2005) X-Ray Diffraction Studies of Amyloid Structure. In Amyloid Proteins. Humana Press, New Jersey. pp. 067–080 http://link.springer.com/10.1385/1-59259-874-9:067. Accessed August 30, 2017.

Makin, O.S., and Serpell, L.C. (2005) Structures for amyloid fibrils. FEBS J 272: 5950–61 http://www.ncbi.nlm.nih.gov/pubmed/16302960. Accessed July 17, 2015.

Margittai, M., and Langen, R. (2008) Fibrils with parallel in-register structure constitute a major class of amyloid fibrils: molecular insights from electron paramagnetic resonance spectroscopy. Q Rev Biophys 41: 265 http://www.ncbi.nlm.nih.gov/pubmed/19079806. Accessed September 5, 2017.

Margot, P., and Karamata, D. (1996) The wprA gene of Bacillus subtilis 168, expressed during exponential growth, encodes a cell-wall-associated protease. Microbiology 142: 3437–3444 http://mic.microbiologyresearch.org/content/journal/micro/10.1099/13500872-142-12-3437. Accessed March 1, 2018.

Michna, R.H., Zhu, B., Mäder, U., and Stülke, J. (2016) SubtiWiki 2.0–an integrated database for the model organism Bacillus subtilis. Nucleic Acids Res 44: D654–62 http://www.ncbi.nlm.nih.gov/pubmed/26433225. Accessed August 31, 2017.

Micsonai, A., Wien, F., Kernya, L., Lee, Y.-H., Goto, Y., Réfrégiers, M., and Kardos, J. Accurate secondary structure prediction and fold recognition for circular dichroism spectroscopy. http://www.pnas.org/content/112/24/E3095.full.pdf. Accessed August 31, 2017.

Ostrowski, A., Mehert, A., Prescott, A., Kiley, T.B., and Stanley-Wall, N.R. (2011) YuaB functions synergistically with the exopolysaccharide and TasA amyloid fibers to allow biofilm formation by Bacillus subtilis. J Bacteriol 193: 4821–31 http://www.pubmedcentral.nih.gov/articlerender.fcgi?artid=3165672&tool=pmcentrez&rendertype=abstract. Accessed January 28, 2016.

Patrick, J.E., and Kearns, D.B. (2008) MinJ (YvjD) is a topological determinant of cell division in Bacillus subtilis. Mol Microbiol 70: 1166–79 http://www.ncbi.nlm.nih.gov/pubmed/18976281. Accessed May 26, 2016.

Petersen, T.N., Brunak, S., Heijne, G. von, and Nielsen, H. (2011) SignalP 4.0: discriminating signal peptides from transmembrane regions. Nat Methods 8: 785–786 http://www.ncbi.nlm.nih.gov/pubmed/21959131. Accessed May 19, 2017.

Petrlova, J., Hansen, F.C., Plas, M.J.A. van der, Huber, R.G., Mörgelin, M., Malmsten, M., et al. (2017) Aggregation of thrombin-derived C-terminal fragments as a previously undisclosed host defense mechanism. Proc Natl Acad Sci U S A 114: E4213–E4222 http://www.ncbi.nlm.nih.gov/pubmed/28473418. Accessed August 30, 2017.

Romero, D., Aguilar, C., Losick, R., and Kolter, R. (2010) Amyloid fibers provide structural integrity to Bacillus subtilis biofilms. Proc Natl Acad Sci U S A 107: 2230–4 http://www.pubmedcentral.nih.gov/articlerender.fcgi?artid=2836674&tool=pmcentrez&rendertype=abstract. Accessed September 27, 2014.

Romero, D., Vlamakis, H., Losick, R., and Kolter, R. (2011) An accessory protein required for anchoring and assembly of amyloid fibres in B. subtilis biofilms. Mol Microbiol 80: 1155–68 http://www.pubmedcentral.nih.gov/articlerender.fcgi?artid=3103627&tool=pmcentrez&rendertype=abstract. Accessed December 13, 2015.

Romero, D., Vlamakis, H., Losick, R., and Koltera, R. (2014) Functional analysis of the accessory protein TapA in Bacillus subtilis amyloid fiber assembly. J Bacteriol 196: 1505–1513 http://www.ncbi.nlm.nih.gov/pmc/articles/PMC3993358/pdf/zjb1505.pdf.

Roux, D., Cywes-Bentley, C., Zhang, Y.-F., Pons, S., Konkol, M., Kearns, D.B., et al. (2015) Identification of Poly-N-acetylglucosamine as a Major Polysaccharide Component of the Bacillus subtilis Biofilm Matrix. J Biol Chem 290: 19261–19272 http://www.ncbi.nlm.nih.gov/pubmed/26078454. Accessed September 1, 2017.

Rufo, G.A., Sullivan, B.J., Sloma, A., and Pero, J. (1990) Isolation and characterization of a novel extracellular metalloprotease from Bacillus subtilis. J Bacteriol 172: 1019–23 http://www.ncbi.nlm.nih.gov/pubmed/2105290. Accessed March 1, 2018.

Schor, M., Mey, A.S.J.S., Noé, F., and MacPhee, C.E. (2015) Shedding Light on the Dock–Lock Mechanism in Amyloid Fibril Growth Using Markov State Models. J Phys Chem Lett 6: 1076–1081 http://pubs.acs.org/doi/abs/10.1021/acs.jpclett.5b00330. Accessed September 6, 2017.

Seemann, T. (2014) Prokka: rapid prokaryotic genome annotation. Bioinformatics 30: 2068–9 http://www.ncbi.nlm.nih.gov/pubmed/24642063. Accessed July 20, 2016.

Seminara, A., Angelini, T.E., Wilking, J.N., Vlamakis, H., Ebrahim, S., Kolter, R., et al. (2012) Osmotic spreading of Bacillus subtilis biofilms driven by an extracellular matrix. Proc Natl Acad Sci 109: 1116–1121 http://www.ncbi.nlm.nih.gov/pubmed/22232655. Accessed September 1, 2017.

Sen, P., Fatima, S., Ahmad, B., and Khan, R.H. (2009) Interactions of thioflavin T with serum albumins: Spectroscopic analyses. Spectrochim Acta Part A Mol Biomol Spectrosc 74: 94–99 http://www.ncbi.nlm.nih.gov/pubmed/19502106. Accessed August 30, 2017.

Serra, D.O., Richter, A.M., and Hengge, R. (2013) Cellulose as an architectural element in spatially structured Escherichia coli biofilms. J Bacteriol 195: 5540–54 http://www.ncbi.nlm.nih.gov/pubmed/24097954. Accessed August 30, 2017.

Serrano, M., Zilhão, R., Ricca, E., Ozin, A.J., Moran, C.P., Henriques, A.O., and Henriques, A.O. (1999) A Bacillus subtilis secreted protein with a role in endospore coat assembly and function. J Bacteriol 181: 3632–43 http://www.ncbi.nlm.nih.gov/pubmed/10368135. Accessed January 14, 2018.

Shewmaker, F., McGlinchey, R.P., Thurber, K.R., McPhie, P., Dyda, F., Tycko, R., and Wickner, R.B. (2009) The Functional Curli Amyloid Is Not Based on In-register Parallel β-Sheet Structure. J Biol Chem 284: 25065–25076 http://www.ncbi.nlm.nih.gov/pubmed/19574225. Accessed August 30, 2017.

Sievers, F., Wilm, A., Dineen, D., Gibson, T.J., Karplus, K., Li, W., et al. (2011) Fast, scalable generation of high-quality protein multiple sequence alignments using Clustal Omega. Mol Syst Biol 7: 539 http://www.ncbi.nlm.nih.gov/pubmed/21988835. Accessed May 19, 2017.

Sipe, J.D., Benson, M.D., Buxbaum, J.N., Ikeda, S., Merlini, G., Saraiva, M.J.M., and Westermark, P. (2016) Amyloid fibril proteins and amyloidosis: chemical identification and clinical classification International Society of Amyloidosis 2016 Nomenclature Guidelines. Amyloid 23: 209–213 http://www.ncbi.nlm.nih.gov/pubmed/27884064. Accessed September 5, 2017.

Sloma, A., Ally, A., Ally, D., and Pero, J. (1988) Gene encoding a minor extracellular protease in Bacillus subtilis. J Bacteriol 170: 5557–63 http://www.ncbi.nlm.nih.gov/pubmed/3142851. Accessed March 1, 2018.

Sloma, A., Rufo, G.A., Rudolph, C.F., Sullivan, B.J., Theriault, K.A., and Pero, J. (1990) Bacillopeptidase F of Bacillus subtilis: purification of the protein and cloning of the gene. J Bacteriol 172: 1470–7 http://www.ncbi.nlm.nih.gov/pubmed/2106512. Accessed March 1, 2018.

Sloma, A., Rufo, G.A., Theriault, K.A., Dwyer, M., Wilson, S.W., and Pero, J. (1991) Cloning and characterization of the gene for an additional extracellular serine protease of Bacillus subtilis. J Bacteriol 173: 6889–95 http://www.ncbi.nlm.nih.gov/pubmed/1938892. Accessed March 1, 2018.

Stöver, A.G., and Driks, A. (1999) Secretion, localization, and antibacterial activity of TasA, a Bacillus subtilis spore-associated protein. J Bacteriol 181: 1664–72 http://www.pubmedcentral.nih.gov/articlerender.fcgi?artid=93559&tool=pmcentrez&rendertype=abstract. Accessed November 30, 2014.

Studier, F.W. (2005) Protein production by auto-induction in high density shaking cultures. Protein Expr Purif 41: 207–34 http://www.ncbi.nlm.nih.gov/pubmed/15915565. Accessed November 6, 2014.

Studier, F.W., and Moffatt, B.A. (1986) Use of bacteriophage T7 RNA polymerase to direct selective high-level expression of cloned genes. J Mol Biol 189: 113–30 http://www.ncbi.nlm.nih.gov/pubmed/3537305. Accessed March 1, 2018.

Sumner Makin, O., Sikorski, P., Serpell, L.C., IUCr, C., D.R., G., D., et al. (2007) *CLEARER* : a new tool for the analysis of X-ray fibre diffraction patterns and diffraction simulation from atomic structural models. J Appl Crystallogr 40: 966–972 http://scripts.iucr.org/cgi-bin/paper?S0021889807034681. Accessed September 13, 2017.

Sunde, M., and Blake, C. (1997) The Structure of Amyloid Fibrils by Electron Microscopy and X-Ray Diffraction. In Advances in Protein Chemistry Volume 50. Elsevier, pp. 123–159 http://linkinghub.elsevier.com/retrieve/pii/S0065323308603204. Accessed August 11, 2017.

Terra, R., Stanley-Wall, N.R., Cao, G., and Lazazzera, B.A. (2012) Identification of Bacillus subtilis SipW as a bifunctional signal peptidase that controls surface-adhered biofilm formation. J Bacteriol 194: 2781–90 http://jb.asm.org/content/194/11/2781.full. Accessed January 7, 2015.

Tran, L., Wu, X.C., and Wong, S.L. (1991) Cloning and expression of a novel protease gene encoding an extracellular neutral protease from Bacillus subtilis. J Bacteriol 173: 6364–72 http://www.ncbi.nlm.nih.gov/pubmed/1917867. Accessed March 1, 2018.

Uversky, V.N., Li, J., and Fink, A.L. (2001) Evidence for a partially folded intermediate in alpha-synuclein fibril formation. J Biol Chem 276: 10737–44 http://www.ncbi.nlm.nih.gov/pubmed/11152691. Accessed September 5, 2017.

Verhamme, D.T., Kiley, T.B., and Stanley-Wall, N.R. (2007) DegU co-ordinates multicellular behaviour exhibited by Bacillus subtilis. Mol Microbiol 65: 554–68 http://www.ncbi.nlm.nih.gov/pubmed/17590234. Accessed July 17, 2015.

Vidakovic, L., Singh, P.K., Hartmann, R., Nadell, C.D., and Drescher, K. (2018) Dynamic biofilm architecture confers individual and collective mechanisms of viral protection. Nat Microbiol 3: 26–31 http://www.nature.com/articles/s41564-017-0050-1. Accessed January 10, 2018.

Vlamakis, H., Aguilar, C., Losick, R., and Kolter, R. (2008) Control of cell fate by the formation of an architecturally complex bacterial community. Genes Dev 22: 945–53 http://www.ncbi.nlm.nih.gov/pubmed/18381896. Accessed September 7, 2017.

Vlamakis, H., Chai, Y., Beauregard, P., Losick, R., and Kolter, R. (2013) Sticking together: building a biofilm the Bacillus subtilis way. Nat Rev Microbiol 11: 157–68 http://www.pubmedcentral.nih.gov/articlerender.fcgi?artid=3936787&tool=pmcentrez&rendertype=abstract. Accessed July 11, 2014.

Whelan, S., and Goldman, N. (2001) A general empirical model of protein evolution derived from multiple protein families using a maximum-likelihood approach. Mol Biol Evol 18: 691–9 http://www.ncbi.nlm.nih.gov/pubmed/11319253. Accessed May 19, 2017.

Wu, X.C., Nathoo, S., Pang, A.S., Carne, T., and Wong, S.L. (1990) Cloning, genetic organization, and characterization of a structural gene encoding bacillopeptidase F from Bacillus subtilis. J Biol Chem 265: 6845–50 http://www.ncbi.nlm.nih.gov/pubmed/2108961. Accessed March 1, 2018.

Yamaguchi, T., Yagi, H., Goto, Y., Matsuzaki, K., and Hoshino, M. (2010) A Disulfide-Linked Amyloid-β Peptide Dimer Forms a Protofibril-like Oligomer through a Distinct Pathway from Amyloid Fibril Formation. Biochemistry 49: 7100–7107 http://pubs.acs.org/doi/abs/10.1021/bi100583x. Accessed August 30, 2017.

Yang, M., Dutta, C., and Tiwari, A. (2015) Disulfide-Bond Scrambling Promotes Amorphous Aggregates in Lysozyme and Bovine Serum Albumin. J Phys Chem B 119: 3969–3981 http://pubs.acs.org/doi/abs/10.1021/acs.jpcb.5b00144. Accessed August 30, 2017.

